# Active regulatory elements recruit cohesin to establish cell-specific chromatin domains

**DOI:** 10.1101/2023.10.13.562171

**Authors:** Emily Georgiades, Caroline L. Harrold, Nigel Roberts, Mira Kassouf, Simone G. Riva, Edward Sanders, Helena S. Francis, Joseph Blayney, A. Marieke Oudelaar, Thomas A. Milne, Douglas R. Higgs, Jim Hughes

## Abstract

As the structure of the genome is analysed at ever increasing resolution it is becoming clear that there is considerable variation in the 3D chromatin architecture across different cell types. It has been proposed that this may, in part, be due to increased recruitment of cohesin to activated cis-elements (enhancers and promoters) leading to cell-type specific loop extrusion underlying the formation of new subTADs. Here we show that cohesin correlates well with the presence of active enhancers and this varies in an allele-specific manner with the presence or absence of polymorphic enhancers which vary from one individual to another. Using the alpha globin cluster as a model, we show that when all enhancers are removed, peaks of cohesin disappear from these regions and the erythroid specific subTAD is no longer formed. Re-insertion of the major alpha globin enhancer (R2) is associated with the appearance of a new peak of cohesin at the site of insertion. In complementary experiments insertion of R2 into a “neutral” region of the genome recruits cohesin, induces transcription and creates a new large (75kb) erythroid specific domain. Together these findings support the proposal that active enhancers recruit cohesin, stimulate loop extrusion and promote the formation of cell specific subTADs.

## Introduction

During interphase, the mammalian genome is organised into a series of hierarchical structures ranging from chromosome territories to smaller domains including active (A) compartments, inactive (B) compartments, lamina associated domains (LADs) and topologically associating domains (TADs). While each of these was originally described at relatively low resolution and in a limited number of cell types, with technological improvements in resolution and by studying wider ranges of cell types, it is clear that there are further levels of organisation. All of these structures are highly dynamic and at high resolution they vary to different degrees, in a cell-type specific manner. For example, although the structure of TADs was originally considered to be relatively constant across cell types, at higher resolution it has become clear that they contain subTADs many of which vary during differentiation, development and cell cycle stage^1–5^. At all stages, a key question has focussed on the relationship between genome structure and function: does structure orchestrate function or is structure a reflection of function?

Since their discovery, the nature of TADs and the mechanisms by which they form has been studied intensively. It is now generally accepted that TADs are formed by the process of cohesin mediated loop extrusion^6–9^. Once recruited, cohesin translocates across chromatin until it reaches a CTCF boundary element in which the N-terminus of CTCF is orientated towards the translocating molecule of cohesin, at which point further extrusion is blocked and cohesin accumulates before it is off loaded by WAPL^10–13^. Cohesin is a multi-subunit protein complex which does not have a specific DNA binding motif, instead current evidence suggests that cohesin is loaded onto chromatin facilitated by the NIPBL-MAU2 complex^14,15^, although there is some debate about the location of cohesin loading. Extensive studies removing the key players (Cohesin, NIPBL, WAPL, CTCF) are consistent with a model in which TADs result from recruitment of cohesin across the entire genome, delimited by the distribution of CTCF sites^16^. Much less is known about how cell-type specific subTADs form and the role that they play in mediating enhancer-promoter interactions. There is building evidence for the recruitment of cohesin via its interaction with NIPBL at activated enhancers and promoters^17–24^, which would go some way to providing a mechanism for cell-type specific domain formation.

It has previously been noted that cohesin is enriched at enhancers within activated subTADs^18,25–27^. To address the specificity of such peaks, we have identified examples of single nucleotide polymorphisms (SNPs) in healthy individuals that either create or abolish an enhancer, which we refer to as ‘natural enhancer variation’. These examples of natural variation have enabled us to determine if there is a correlation between cohesin occupancy specifically at these regions with otherwise minimal perturbation of the locus.

We utilised the well characterised mouse alpha globin locus to ask whether enhancer elements may recruit cohesin and thereby initiate the formation of a cell-specific 3D domain bringing enhancers and promoters into close proximity. We have previously shown that the alpha-cluster is contained within a 165kb TAD found in all cell types tested, whereas a 65kb evolutionarily conserved, erythroid-specific subTAD containing all alpha-like genes and five erythroid-specific enhancers (R) in the order 5’ R1-R2-R3-Rm-R4-ζ-α1-81-α2-θ2-3’ forms during erythroid differentiation. The subTAD starts to form early in erythroid differentiation prior to alpha globin gene expression, at which time the cis-regulatory elements are bound by erythroid specific transcription factors (TFs)^28,29^. By the end of differentiation, the enhancers and promoters are in close proximity and the alpha globin genes are highly transcribed. Of interest, we have previously shown that there is enrichment of both cohesin and NIPBL at the alpha globin enhancers in erythroid cells^17,30^, and that inserting CTCF sites between the enhancers and promoters has a predominant effect on alpha globin expression when the N-terminus of these sites prevents translocation from the enhancers to the promoters^31,32^. This suggested that these cis-regulatory elements play an important part in the formation of the subTAD^33,34^ however given the complexity of this relatively simple locus further experiments described here are required to test this hypothesis.

In this study we aimed to determine if active enhancers were sites of cohesin recruitment. To do this, we took advantage of natural variation in human sequences and identified unique enhancers in different individuals and showed that cohesin displayed distinct recruitment patterns that reflected the presence of unique active enhancers. Further, we took advantage of the well characterized globin locus to create an engineered artificial locus to ask whether a single classical enhancer (R2) rather than the complex super-enhancer (R1, R2, R3, Rm, R4) is capable of forming a sub-domain. Third, to remove all confounding elements and environment, we identified a “neutral” region of the genome and inserted R2 on its own to determine if it could recruit cohesin and create an erythroid-specific chromosomal domain independently. Together these experiments strongly suggest that R2 on its own is able to recruit cohesin and form an erythroid specific subTAD.

## Results

### Cohesin colocalises with active enhancers

It has been proposed that cohesin is loaded at active regulatory elements across the genome^17–24^. Although to our knowledge this has not been quantified on a genome-wide scale. To look at this in detail, we selected three donors at random from the Oxford Biobank and performed histone modification ChIP-seq for H3K27Ac, H3K4me1 and H3K4me3, along with CTCF ChIP-seq and ATAC-seq in CD34+ erythroid cells to characterise the regulatory landscape in these individuals and classify the regulatory elements. We identified sites at which there was a natural variation in enhancer activity between individuals and defined these sites of ‘natural variation’ to be sites which have open chromatin and are marked with both H3K27Ac and H3K4me1 in at least one, but not all three donors, most likely representing variant enhancers. Then we performed RAD21 ChIP-seq in CD34+ erythroid cells from three donors to determine whether there was a correlation between enhancer activity and cohesin recruitment. Using allelic skew analysis (details in Materials and Methods), we identified 51 robust examples of polymorphic enhancers in which the associated enhancer was either active or inactive. For each of these regions the RAD21 coverage was calculated and plotted to show a positive correlation between RAD21 signal and enhancer activity (Fig. 1A).

**Figure 1A:**
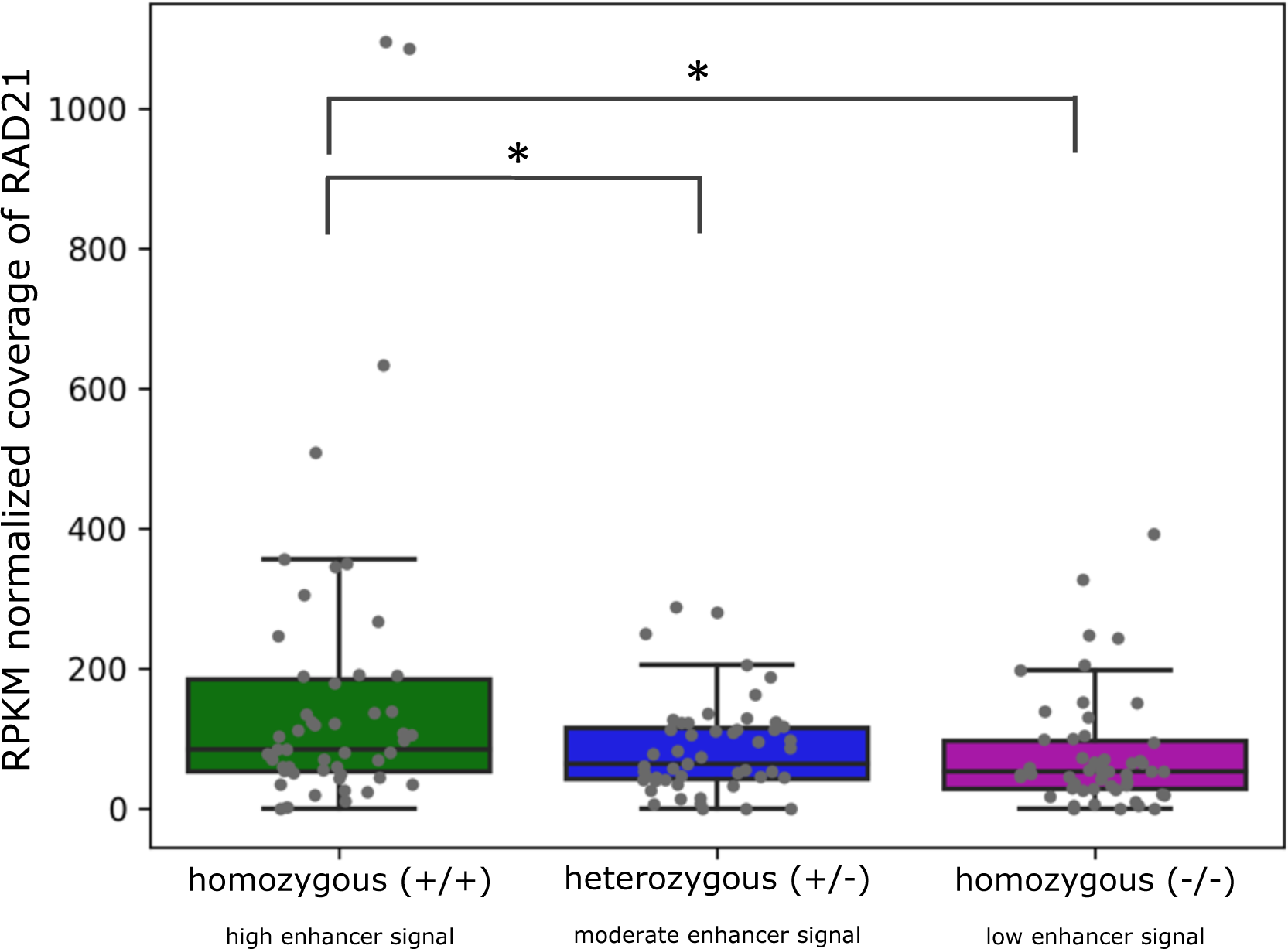
There is a positive correlation between the presence of enhancer signal and coverage of RAD21 suggesting active enhancers accumulate cohesin. For 51 regions in which there is a robust difference in ATAC-seq signal across the donors, RAD21 signal was calculated for the three donors and the donors classified as either homozygous (+/+ or -/-), or heterozygous (+/-) for enhancer signal. Each of the 51 individual data points are plotted as grey dots. Statistical significance between pairs was calculated using the T-test, asterisk indicates p-value ≤ 0.05. Coverage was normalized by RPKM and by region size. For further explanation of analysis refer to Materials and Methods.

Fig. 1B, as well as two additional examples provided in Fig. S1 and S2, show examples of sites included in the meta-analysis in which there is clear enhancer variation between donors, and a correlation between enhancer activity and cohesin occupancy in these cases. To determine the cause of these variations in enhancer activity between donors we used a basic peak calling approach, prioritising sites with enhancer variation. We then used phased genome data to identify allele-specific genetic variation correlated with the presence of an enhancer. Fig.1B shows one example in which we have identified genetic variants correlated with either the gain or loss of an enhancer. Here we have identified haplotypes associated with the gain or loss of cohesin signal. This was achieved using phased genomes to identify any skew in the ATAC signal. In doing so we determined the key variants which segregate these haplotypes and formed our hypothesis of causality. The examples shown here could be seen as ‘experiments of nature’, in which some enhancer elements are present seemingly randomly across the genome and between individuals and providing an additional line of evidence of a link between enhancer activity and cohesin recruitment. In these examples we were unable to disentangle whether the correlation with cohesin occupancy was directly due to the activity of the enhancers, given that we have only studied one cell type and these “putative enhancers” have not been experimentally validated. Therefore, to more rigorously investigate this hypothesis, we sought out a highly characterised locus in which the enhancer elements could be studied in a both active and inactive cell types.

**Figure 1B:**
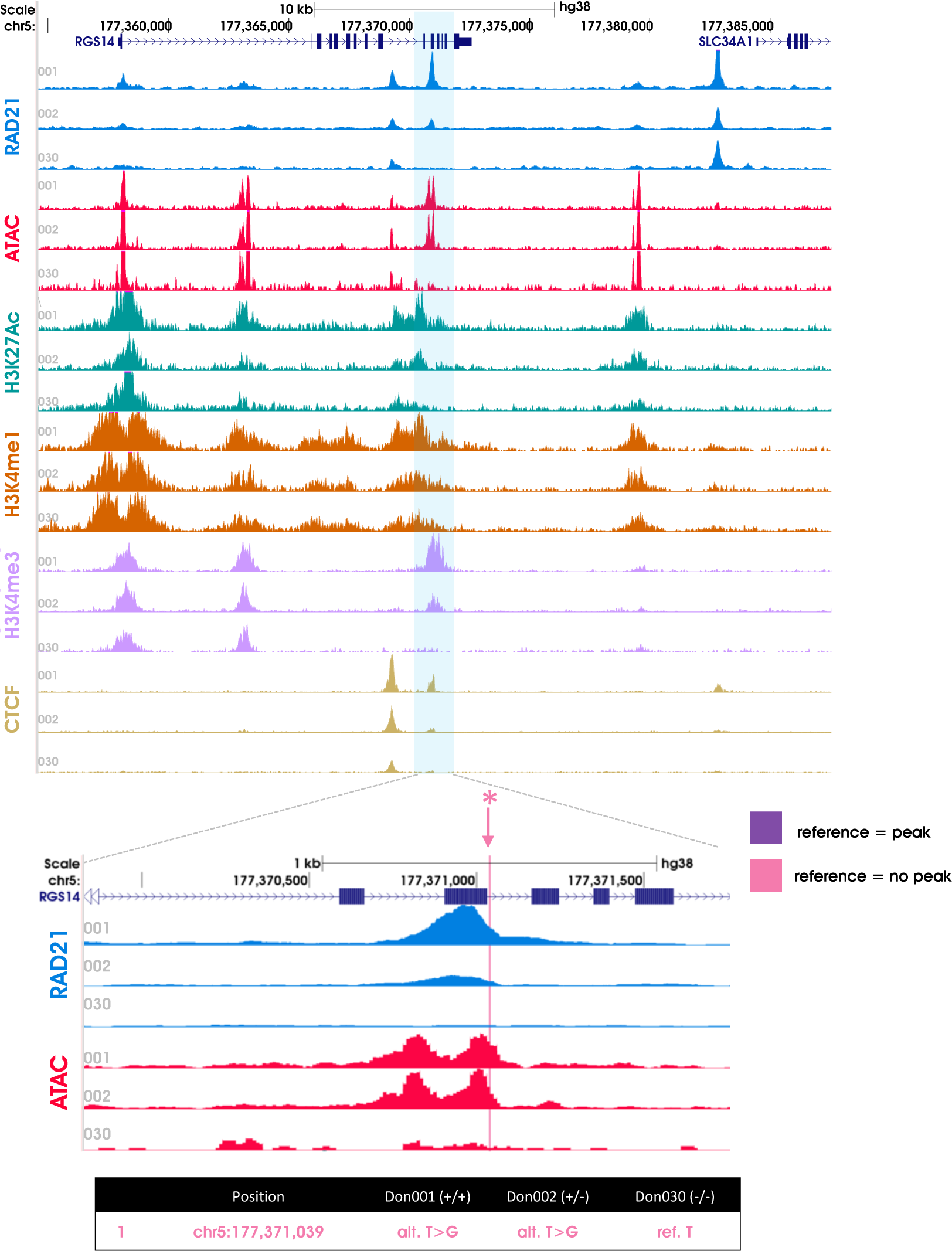
An example of a site with natural variation of enhancer signal across the three donors and the correlation with cohesin occupancy. Region shown is chr5:177,367,955-177,373,964 (hg38). Numbers 01, 02, 30 indicate the anonymized donor identifiers. ChIP-seq for RAD21, CTCF and histone marks (H3K4me1, H3K4me3, H3K27ac) are shown for each donor in the top panel along with the open chromatin signal (ATAC-seq). The region highlighted in blue is shown in detail below. Here the difference in signal across the donors can be clearly seen: donor 01 is homozygous (+/+) and displays a positive ATAC-seq signal and the greatest peak of RAD21, donor 02 is heterozygous (+/-) and displays a positive ATAC-seq signal and a moderate peak of RAD21, donor 30 is homozygous (-/-) with no ATAC-seq or RAD21 signal. The pink line indicates the SNP we have identified as potential causal for this difference across the donors. Two additional examples are provided in Fig. S1 and S2.

### The alpha globin super-enhancer is a useful model in which to controllably test cohesin recruitment to active enhancers

The clusters of both human and mouse alpha globin enhancers (R1-R4) fulfil the definition of super-enhancers^34–36^. This locus is not only one of the best studied regulatory regions of the genome, there are also a number of robust cellular models in which to study the locus in both its active (erythroid) and inactive (non-erythroid) states. To study the recruitment of cohesin to these enhancer elements, we made use of two previously described models: a mouse embryonic stem cell (mESC) line in which all of the five enhancers have been deleted (super-enhancer knock out (SE KO)^33^) and another model in which all enhancer elements except for the strongest of the five enhancers (R2) have been deleted (R2-only^33^). To study the enhancer activity in these models, CD71+ erythroid cells from embryoid body cultures^37^ and Ter119+ primary erythroid cells from mice were obtained to study the SE KO and R2-only models respectively. RAD21 ChIP-seq was then performed on the SE KO, R2-only and WT control CD71+ erythroid cells. Peaks of cohesin were found in the WT cells at all of the five enhancer elements (Fig. 2A). By contrast, in the SE KO model, no peaks of cohesin were found at any of the enhancer elements. Importantly, in the R2-only model cohesin was only observed at the site of R2 enhancer element. In both models, similar peaks of cohesin at the CTCF sites could be seen as those observed in WT cells. By visual inspection, it appeared that between the two convergent CTCF sites containing the alpha globin subTAD there is a greater build-up of cohesin across the domain in the WT compared to both enhancer KO models (Fig. 2A). To quantify this observation, the total ChIP’d sequence coverage of RAD21 across the region between the convergently orientated CTCF sites flanking the subTAD was compared with a nearby size-matched region (∼61.5kb) outside of the CTCF sites and a ratio calculated. Fig. 2B shows that cohesin within the SE region is enriched ∼6 fold in the WT compared to the enhancer KO models, this shows that in the presence of all 5 alpha globin enhancers, recruitment of cohesin across the entire subTAD is substantially increased. When these enhancers are knocked out, cohesin is not recruited into the subTAD to the degree normally observed in WT erythroid cells.

**Figure 2A:**
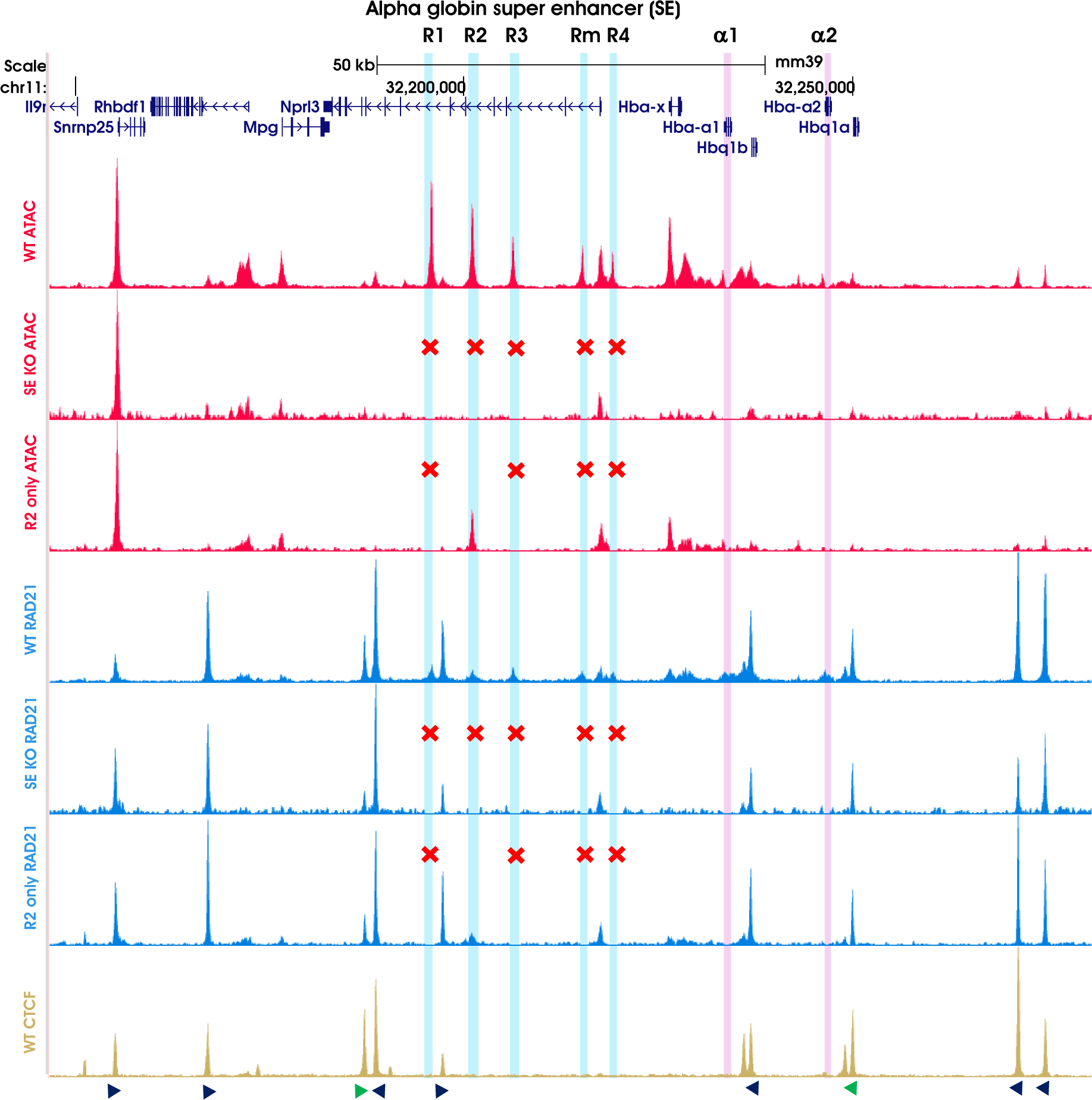
Cohesin accumulates at the sites of active enhancers. The region spanning the alpha globin locus (chr11:32,146,649-32,280,800, mm39) is shown. Alpha globin enhancers are highlighted in blue and the two alpha genes (a1 and a2) highlighted in pink. GENCODE VM32 genes within the region are shown at the top in dark blue. ATAC-seq and ChIP-seq tracks for RAD21 and CTCF are shown with corresponding model indicated to the left. Red crosses above tracks indicate which of the enhancers are knocked out. All bigwig tracks are RPKM normalised and plotted with bin size 1. The orientation of CTCF motifs are shown below the CTCF track, the two highlighting in green indicated the boundaries for the region analysed for Figure 2B.

**Figure 2B:**
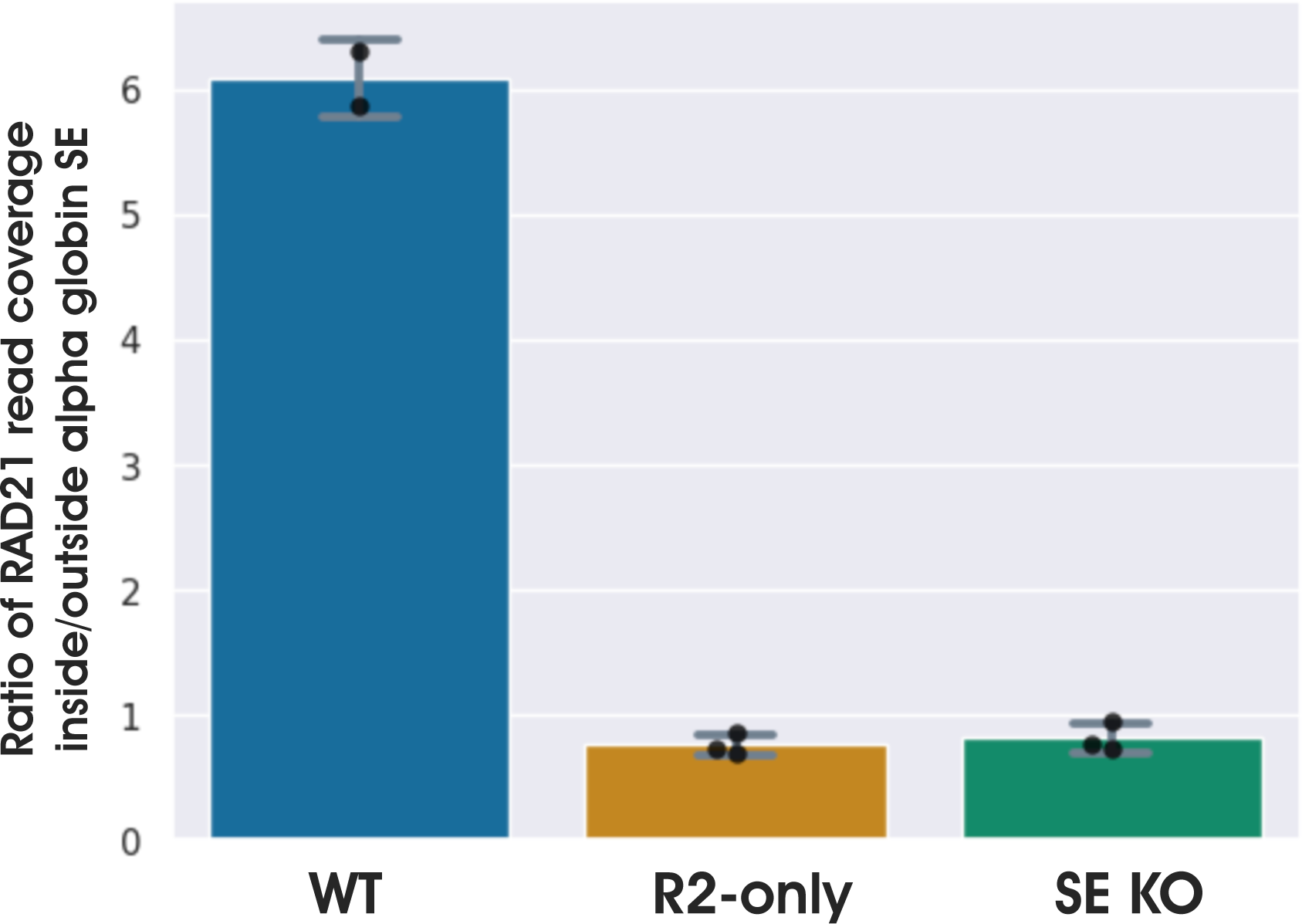
Cohesin residency within the region of the alpha globin SE is significantly reduced upon deletion of the enhancer elements. Bar plot showing the ratio of cohesin coverage inside (chr11:32,188,452-32,249,902) versus outside (chr11:32,300,054-32,391,239) the alpha globin SE region. Two replicates for WT and three replicates for the R2-only and SE KO model are plotted; each point represents one replicate. Error bars represent standard deviation across replicates.

### Active enhancers give rise to tissue-specific subTADs

Cohesin is known to play an important role in the establishment of 3D genome structure through its role in loop extrusion^38–41^. Since fully or partially deleting the SE changes the amount of cohesin recruited to the active alpha globin subTAD in erythroid cells we addressed how this might impact the 3D genome structure of this locus through the interaction between the enhancers and their cognate promoters. First, we performed Tiled-C in undifferentiated mESCs in which the alpha globin locus is inactive, and genes are not transcribed, and compared this to a fully activated state in WT erythroid cells derived from mouse fetal liver (Fig. 3A). When plotted as a log2 comparison plot, changes in interaction frequencies across the alpha globin locus can be readily observed. Next, we looked at how these patterns might change by comparing R2-only erythroid cells^33^ with inactive mESCs (Fig. 3B). This showed a pronounced difference in the interaction frequencies around the alpha globin locus in the R2-only erythroid model compared to the inactive mESCs. Finally, a comparison of the interaction frequencies of WT erythroid cells and the R2-only erythroid cells was plotted (Fig. 3C), which demonstrated the extent to which the activity of the additional enhancers in the WT cells contribute to a greater degree of interactions compared to when only one of the enhancers is present. These data demonstrate that even a single active enhancer can direct the recruitment of cohesin to the alpha globin cluster and influence its 3D structure via loop extrusion albeit with a reduced capacity than when all native enhancers are active. We conclude that the number of active enhancers controls the recruitment of cohesin within the domain and directly affects the degree of 3D interactions.

**Figure 3:**
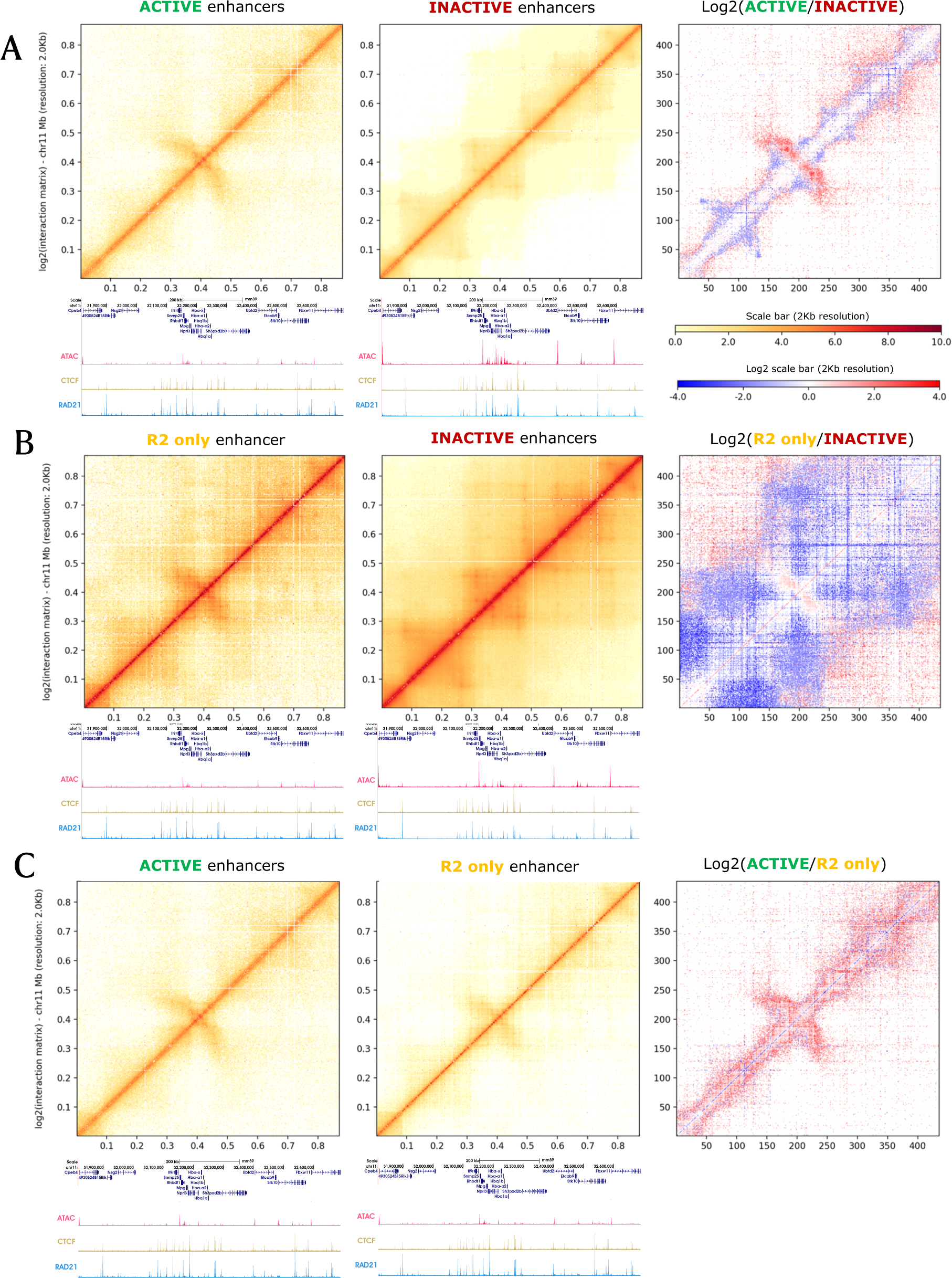
Activation of erythroid specific alpha globin enhancers coincides with a change in 3D domain structure. Tiled-C heatmaps across region chr11:31,818,001-32,690,000 (mm39) at 2kb resolution. Tiled-C in WT mESC was generated for inactive enhancers, in WT fetal liver for active enhancers and R2 only fetal liver model for R2 only. (A) Alpha globin enhancers are inactive in ES cells, and active in Ter119+ fetal liver cells. Data from the two states were normalized before plotting a log2 comparison of the chromatin interactions across the locus in an inactive versus active states. ATAC and ChIP-seq tracks are from the corresponding models. (B) The inactive alpha globin locus in ES cells is next compared to the model in which only the R2 enhancer is present and active. Data from the two states were normalized before plotting a log2 comparison of the chromatin interactions across the locus in an inactive versus R2 only active states. ATAC and RAD21 ChIP-seq tracks are from the corresponding models, CTCF ChIP-seq is from WT mESC for the inactive model and WT embryoid body erythroid cells for the R2 only model. (C) The active alpha globin locus in Ter119+ fetal liver cells is next compared to the model in which only the R2 enhancer is present and active. Data from the two states were normalized before plotting a log2 comparison of the chromatin interactions across the locus in a fully active versus R2 only active states. ATAC and ChIP-seq tracks are from the corresponding models. Note: Datasets have been normalized in a pairwise manner for each A, B and C to account for the differences in sequencing depth across experiments, therefore the individual plots appear different across A, B and C.

### Identification of a neutral but activatable region of the genome

An important limitation of using the enhancer KO models to investigate the independent role of an enhancer in recruiting cohesin to the cluster is that it is impossible to control for all potentially confounding factors that may contribute to activity of the enhancer. Whilst the R2-only and SE-KO models provide useful insights into how an enhancer might influence cohesin recruitment, the results may be influenced by activity from neighbouring active genes, and/or other regulatory elements within or surrounding the locus. We therefore developed an orthogonal approach to study cohesin recruitment by the R2 enhancer by inserting this element outside of its normal chromatin context, in a region of the genome devoid of gene activity. We therefore needed to identify a suitable region of the genome in which to insert the R2 enhancer to determine if this element is capable of recruiting cohesin in an independent manner.

To increase the efficiency of the genome editing strategy, we chose to use a male mESC line and target the X chromosome as this circumvents the need to isolate homozygously edited clones. The target locus needed to be amenable to activation: therefore, the search was limited to regions within the A compartment of the genome^42^ to avoid inserting into repressive LADs that can restrict chromatin accessibility and TF binding. To classify the A and B regions, compartmentalisation analysis was performed on published Hi-C data in mESCs^43^. Once we had identified an activatable region, we analysed ChIP-seq and ATAC-seq data from Ter119+ primary erythroid cells and mESCs to assess the chromatin landscape. To avoid confounding effects, whilst the region needs to be activatable (i.e., not epigenetically repressed), the ideal region should be as neutral as possible. We therefore looked for a region devoid of transcriptionally repressive marks (H3K9me3 and H3K27me3), open chromatin signal (determined by ATAC-seq), ChIP-seq signals for histone modifications typically associated with active enhancers and promoters (H3K4me1 and H3K4me3 respectively) or annotated genes. Of the candidate regions on chromosome X, one satisfied all criteria for being potentially neutral and activatable chromatin in both mESCs and primary erythroid cells: an approximately 110 kb region at chrX:11,224,970-11,335,361 (mm39) (Fig. S4). To ensure there was no underlying complex chromosome structure other than the normal proximity signal expected of the chromatin fibre in this region, DpnII fragments ranging between 1-3 kb in length were identified across the locus to act as viewpoints of interest, and capture oligonucleotides were designed to six viewpoints across the locus (Table S1, also labelled in Fig. S4 and S5). Capture-C was then performed across this locus in wild type mESCs and CD71+ erythroid cells isolated from embryoid body cultures, this generated interaction data of a higher resolution than the available Hi-C data and confirmed the absence of any recognisable structure in the two cell types (Fig. S5). The DpnII viewpoint fragments were also used as ‘landing pads’ (LPs) in which to target the R2 alpha globin enhancer element, giving the ability to assay local chromatin structure pre- and post-editing with the same set of capture oligonucleotides.

### Inserting the R2 enhancer into a neutral region of the genome results in the formation of a new erythroid-specific domain structure

A 325bp segment of DNA containing the sequence for the R2 alpha globin enhancer was targeted to the second landing pad (LP2) in the chrX locus in mESCs using CRISPR-Cas9-mediated homology directed repair (HDR). This edited cell line is referred to as ‘R2-insertion’ model. To determine whether the inserted R2 became accessible and caused changes in local chromatin in ESCs, ATAC-seq was performed on R2-insertion and WT ESCs. R2 is a known erythroid-specific enhancer, therefore in undifferentiated ESCs, in which the enhancer is inactive, no change in accessibility was expected or observed. R2-insertion and WT mESCs were then differentiated to form embryoid bodies and CD71+ erythroid cells isolated at day seven of differentiation, at which point the R2 enhancer is expected to be active, and ATAC-seq performed.

ATAC-seq reads from this cell line were aligned to a custom reference genome allowing for multi-mapping of reads that mapped to both the native R2 and the inserted R2 sequence. Reads that originated directly from the 325 bp R2 sequence were randomly mapped to one of the two genomic locations harbouring R2 resulting in peaks at both locations. To confirm that the accessible chromatin peak over the inserted R2 was a true signal, rather than a false-positive signal originating from native R2, the profiles of the paired-end reads spanning the junctions of the R2 insertion were compared to the junctions at native R2. A higher total count of R2 reads than WT was shown in all three clones (Fig. S3). Interestingly, the activation of the R2-insertion cells resulted in both local changes in accessibility in the vicinity of the insertion on chromosome X, and the generation of novel ATAC-seq peaks up to ∼75 kb downstream of the insertion site (Fig. 4A). These results show that the inserted R2 enhancer element exerts changes on surrounding chromatin over a substantial distance suggesting the establishment of long-range interactions upon its insertion. Together these findings show that the ectopically inserted R2 sequence acts as a long-range cis-regulatory element.

**Figure 4A:**
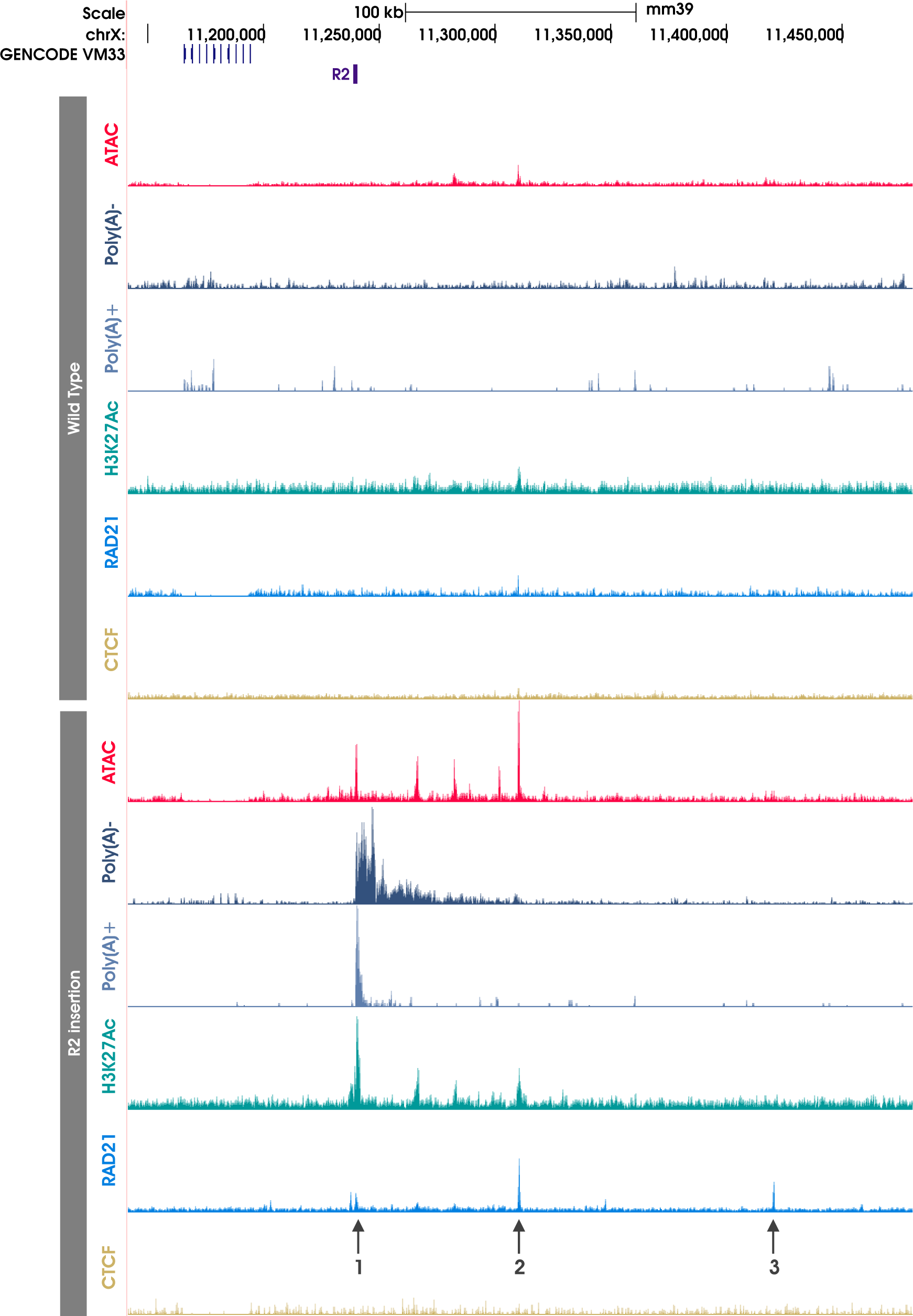
Insertion of the R2 enhancer element gives rise to novel peaks of open chromatin, H3K27Ac, RAD21 and *de novo* transcription. All data is generated in CD71+ erythroid material derived from embryoid body culture. Top panel shows the data generated in WT cells; bottom panel shows data from R2-insertion model cells. Arrows highlight the RAD21 peaks discussed in the text. The location of the R2 insertion is shown as a purple rectangle. A putative CTCF motif identified to overlap RAD21 peak 3 (show at arrow 3) by FIMO analysis, however no signal seen by CTCF ChIP-seq. All data is RPKM normalised and aligned to the mm39 reference genome.

H3K27ac is associated with active chromatin and marks regions of active transcription, H3K4me3 is associated with active promoters and H3K4me1 is associated with active, or primed enhancers (although it is also found at active promoters). ChIP-seq for the histone marks H3K27ac, H3K4me1, and H3K4me3, were performed in the R2-insertion model and WT erythroid cells to characterise any changes in the regulatory landscape following the insertion of R2 (Fig. S6 and S7). The novel ATAC-seq peaks appearing downstream of the R2-insertion in erythroid cells were marked with H3K27ac (Fig. 4A) and H3K4me1 (Fig. S7). Interestingly, unlike the downstream elements, the inserted R2 sequence itself was also marked with H3K4me3 in erythroid cells (Fig. S7). This was unexpected given the histone marks observed at R2 in the native alpha globin environment (Fig. S6B). Therefore, unexpectedly, the inserted R2 enhancer had the chromatin signature of an active promoter whereas the newly formed accessible regions of chromatin lying downstream of the inserted R2 had the signatures of enhancers.

Initially, characterisation of the chromosome X region was performed in primary erythroid cells derived from fetal liver. However, in order to make a direct comparison between an active and inactive environment, we performed the R2-insertion experiments in erythroid cells derived from mESCs so we could assay the inactive (mESC) and active state (embryoid body derived erythroid cells), something that is not possible when using primary material. A caveat of using erythroid cells derived from different sources was the possibility that that the chromatin in embryoid body derived erythroid cells differed from that in the fetal liver derived cells. Indeed, in the WT mESCs we observed very small pre-existing peaks of H3K4me1 corresponding to the peaks which arise to a much greater extent when R2 is inserted upstream (Fig. 4A and Fig. S7). The peaks which appear downstream upon the insertion of the R2 element mark are marked by low-level H3K4me1 in the unedited cells. This suggests that there is some low-level cryptic activation at these sites supported by the existence of erythroid specific TF motifs such as Gata1 beneath these sites. This cryptic activity is then increased under the influence of the inserted active R2 element.

Given that the newly inserted R2 enhancer element in R2-insertion erythroid cells had the signature of an active promoter, we asked whether this initiated RNA transcription. Strand specific Poly(A)- and Poly(A)+ RNA-seq was performed on R2-insertion and WT erythroid cells (Fig. 4A). In both Poly(A)- and Poly(A)+ data, unidirectional transcription was observed. In Poly(A)-, transcription was seen from the R2 insertion site, producing a ∼75kb transcript including the regions containing the novel enhancer-like elements; and in Poly(A)+, a ∼4 kb transcript from the inserted R2 was observed. Together, these data show that the inserted R2 in the R2-insertion model acts as a promoter-like regulatory element producing both non-polyadenylated and polyadenylated, unidirectional transcripts and activating cryptic elements over a region of 75kb.

### The α-globin R2 enhancer sequence is sufficient to form a regulatory domain in erythroid cells

To investigate whether the insertion of R2 altered the 3D chromosome conformation in and around the insertion site, Capture-C was performed from the viewpoint of LP2 (R2 insertion site) in both R2-insertion and WT erythroid cells. As expected, in WT erythroid cells Capture-C showed a normal distribution of interactions around the viewpoint, consistent with the proximity signal of a chromatin fibre with no underlying regulatory structure (Fig. 4B). By contrast, in the R2-insertion erythroid cells there were new interactions between the inserted R2, and the region downstream (3’) extending for 75kb including the novel enhancer-like elements. In addition, a slight enrichment in interactions upstream (5’) of LP2 in R2-insertion erythroid cells was observed, although these interactions did not extend as far as those downstream.

**Figure 4B:**
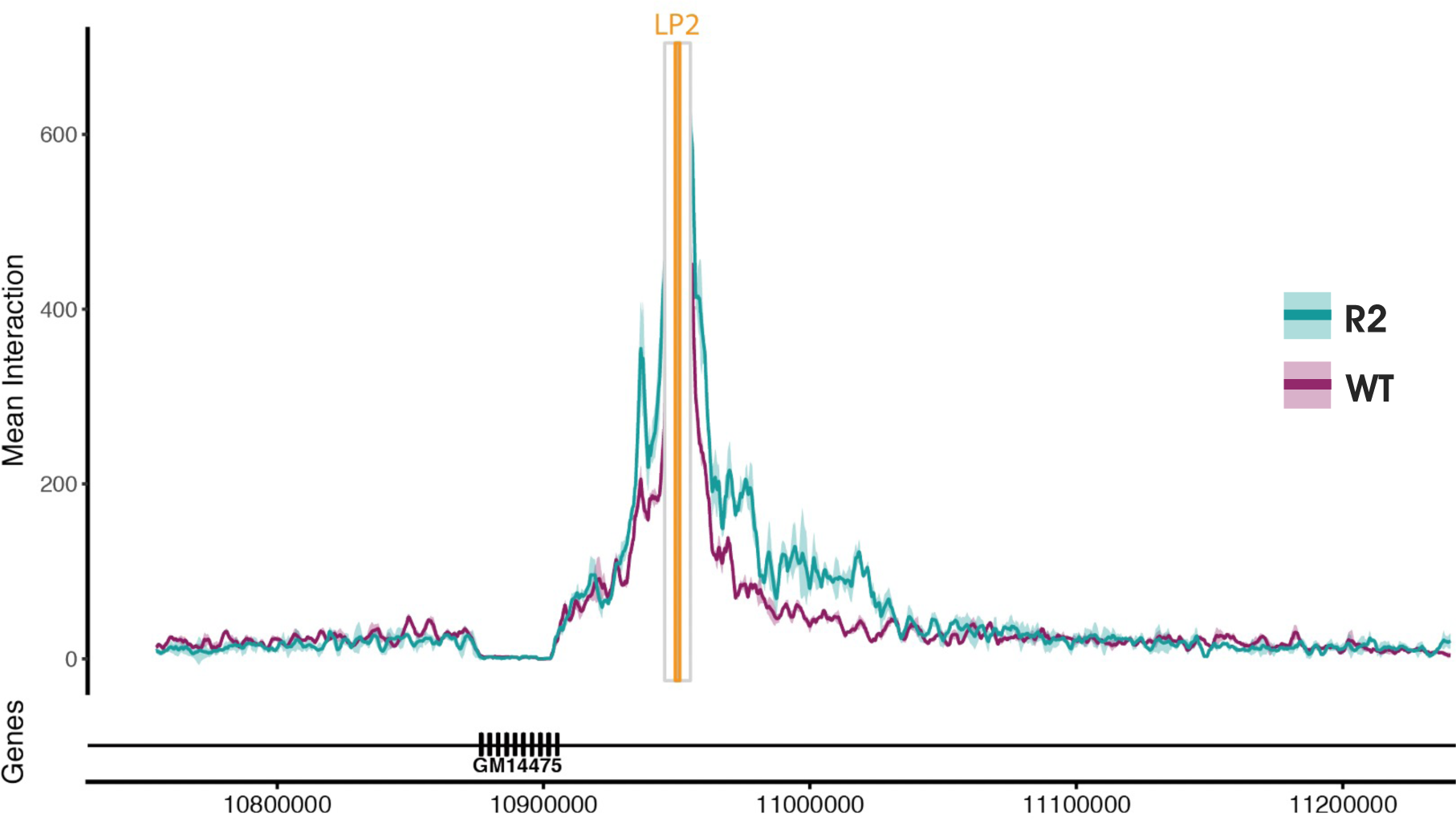
The inserted R2 element contacts sites upstream and downstream suggestive of domain formation. Comparison of the local genome topology around the R2 insertion site in R2-insertion erythroid cells versus WT. NG Capture-C interaction profile from the R2 (orange line) with a 1kb exclusion zone around the viewpoint. WT profile is shown in purple and R2-insertion in turquoise. Each profile represents normalised, averaged unique interactions from three biological replicates. Standard deviation is represented as a halo around the average across a 3kb sliding window. Genomic location and relative positioning of genes is shown below the interaction profile (mm9).

Capture-C interactions were not specifically focused at the novel enhancer-like elements, but rather extended over the entire domain encompassing the region of active transcription. These findings suggest that insertion of the R2 element might promote the formation of a new cell-specific chromatin domain, however only one-way interactions from one viewpoint (LP2) have been characterised and therefore could not be used to make inferences about higher-order structures and configurations. To investigate the 3D interactions across the whole of the locus in an unbiased manner, Tiled-C was performed across a ∼3.3Mb region spanning the inserted R2 in the R2-insertion and in WT erythroid cells (Fig. 4Ci and 4Cii). In agreement with the Capture-C data, Tiled-C data revealed an increase in the interaction frequencies of the R2 insertion site with the surrounding locus in the edited cells compared to WT, reminiscent of a TAD or subTAD-like structure.

**Figure 4C:**
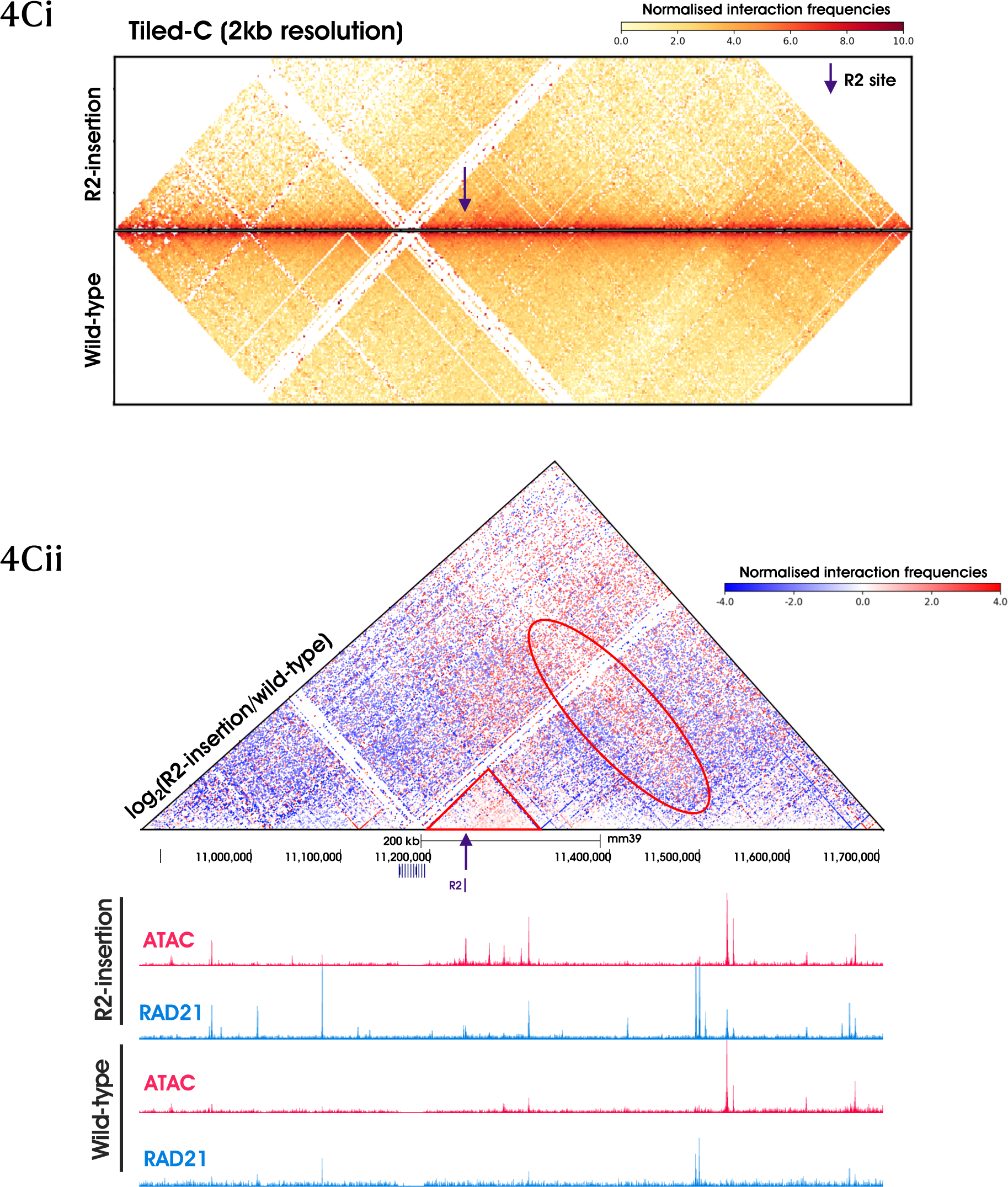
The R2 enhancer can initiate the formation of a subTAD when isolated outside of its usual chromosome context. (i) Tiled-C heatmaps showing normalized interaction frequencies shown at 2kb resolution across region chrX:10,875,000-11,705,000 (mm39). Arrow indicates the site in which R2 is inserted. (ii) A log_2_ comparison plot of the two normalized Tiled-C experiments shown in (i). Triangle contains a region of increased interactions when the R2 insertion model is compared to the WT and indicates the formation of a new subTAD. The oval outline indicates an additional stripe of newly formed interactions which suggest the activated elements within the domain are also forming longer-range interactions beyond the newly formed subTAD. Tandem repeats of the H2al1 genes result in multi mapping issues which explains the lack of signal detection from the region surrounding these genes. ATAC-seq (dark pink) and RAD21^TST^ (blue) ChIP-seq data is shown for reference.

### Activation of the R2 erythroid enhancer is linked with cohesin recruitment

We hypothesised that proteins may be loading at, and translocating from, the newly inserted R2 element and subsequently directing loop extrusion and long-range chromatin interactions. Although transcription per se has been implicated in this process, there is now abundant evidence that the cohesin complex plays the major role in loop extrusion. To analyse the binding of cohesin at this engineered locus, we performed ChIP-seq of the cohesin component RAD21. The endogenous RAD21 protein was tagged with a twin Strep-tag (RAD21^TST^) and ChIP-seq was performed in R2-insertion and WT erythroid cells Fig. 4A (also Fig. 4Cii). A small peak of cohesin is observed at the R2 insertion site in R2-insertion erythroid cells (arrow 1 in Fig. 4A), and this corresponds to the open chromatin site identified through ATAC-seq. This is consistent with the newly inserted R2 element acting as a recruitment site for cohesin as reported for other enhancers and promoters^17–24^.

The process of cohesin-mediated loop extrusion is usually delimited by CTCF binding sites and enriched peaks of cohesin appear at these sites when the translocation of cohesin is stalled by CTCF. Other mechanisms which stall the translocation of cohesin have been reported^24,44,45^. While CTCF ChIP-seq data in both WT erythroid cells and WT mESC did not show any significant peaks in this domain (Fig. 4A) we observed a strong cohesin peak 71kb downstream of the enhancer (arrow 2 in Fig. 4A). This coincided with the previously identified ATAC peak at this site. It could be that the presence of a strong enhancer instates CTCF binding within the vicinity. However, using FIMO analysis, a putative negatively oriented CTCF binding site was found to overlap the cohesin peak at the most downstream open chromatin site ∼182kb downstream of the inserted R2 sequence (chrX:11,420,541-11,420,558, arrow 3 in Fig. 4A). However, using ChIP-seq, no significant peaks of CTCF were identified in this region. Similarly, no CTCF peak was seen at a second ATAC peak downstream of R2 (arrow 2 in Fig. 4A). Such peaks of cohesin not associated with CTCF sites have been previously noted^46^: it has been suggested that a small fraction of cohesin does not colocalise with CTCF and instead resides at sites marked by active chromatin features and bound by cell type specific TFs^18,25,47^, which could be the case in the R2-insertion model.

## Discussion

In this work we have presented three different approaches to studying the recruitment of cohesin to active enhancer elements. It has been recently shown that single nucleotide mutations can give rise to de novo enhancers in brain^48^. We speculated that common variants in the form of SNPs that vary across individuals may be correlated with the presence or absence of enhancer activity and could therefore be used as naturally occurring examples in which to study how cohesin recruitment varies with enhancer activity. Using three individuals from the Oxford Biobank we identified numerous examples of enhancer variation and used these to show a correlation between enhancer activity and cohesin occupancy. This data suggests that the activity of an enhancer somehow influences the recruitment of cohesin. However, in the examples shown there are also differences in the CTCF binding at these enhancers which therefore does not rule out the possibility that there is no difference in cohesin recruitment, but rather the observed pattern is due to the increased boundary effect at the CTCF sites. We therefore looked to use a model in which we could controllably activate the enhancers to compare cohesin recruitment and 3D genome architecture in the inactive versus active state, in a well characterised locus.

We deployed knock out models in which either all, or all but one of the enhancers in the alpha globin super-enhancer were removed, but CTCF sites kept intact, to investigate how cohesin recruitment and the domain structure changed. Interestingly, we showed that in WT erythroid cells there was approximately six times more RAD21 within the region containing the alpha globin super-enhancer than outside the region, whereas in the SE KO and the R2-only model this ratio was reduced to approximately 1:1. This striking result suggests that the presence of the enhancers have some influence on the recruitment of cohesin, however it is curious that the R2-only model and the SE KO show a very similar ratio. If the hypothesis of enhancers recruiting cohesin is correct, it would be expected that the ratio for R2-only would be a fifth of the WT, given that the number of recruitment sites is reduced by 4, and therefore would still show an enrichment compared to the SE KO model. However, it has previously been shown that the expression of the alpha globin genes is reduced to 10% of WT levels in the R2-only model^33^ suggestive of a synergy between the enhancer elements which influence each other within the context of the super-enhancer; the non-linear reduction in cohesin recruitment aligns with this hypothesis.

Next, we isolated the strongest of the alpha globin enhancers in a neutral region of the X chromosome to investigate if this single enhancer was sufficient to recruit cohesin and form a chromatin domain. This was particularly intriguing as the insertion of this short 325bp sequence containing the R2 enhancer was able to initiate both drastic changes in the regulatory landscape and transcription. When we investigated the 3D architecture following the insertion, we observed the formation of a domain-like structure across the region which when correlated with the cohesin ChIP-seq data supports the model of loop extrusion as a mechanism for this domain formation. Whilst we speculate that the inserted enhancer element, once activated in the erythroid lineage, is responsible for the recruitment of cohesin which itself is able to direct domain formation, there are a number of alternative hypotheses. One of the most puzzling observations is the tendency for the R2 element, once inserted into chrX, to exhibit high levels of H3K4me3 and unidirectional transcription, which would ordinarily be indicative of a promoter element. It remains to be understood how this can be the case and provides further weight to the idea that regulatory elements should be considered to have a spectrum of activity rather than being classified into distinct dichotomous groups of either promoters or enhancers^49–51^.

It has been estimated that a high proportion of the genome carries regulatory potential^52^, it is therefore perhaps not surprising that it is challenging, or potentially impossible to find a truly neutral region of the genome. When we first set out to identify a region of the genome in which to insert the enhancer element, the erythroid material we had access to and for which we used to characterise the chrX locus originated from Ter119+ erythroid cells derived from spleen on the assumption that the chromatin environment of mouse erythroid material should be consistent across different sources. We therefore expected that that chromatin marks in the embryoid body material to be identical, which was not the case. Instead of a neutral region devoid of chromatin marks as was the case in the spleen derived cells, the embryoid body derived erythroid material exhibited some minor peaks of H3K4me1 suggestive of cryptic enhancer like elements (Fig. S4). Whilst we did not start from a completely blank genomic canvas by targeting this region in WT cells, we did show that this region was inactive in terms of transcription and 3D structure (Fig. 4A-C), despite the presence of cryptic enhancers. Therefore, when the R2 enhancer was inserted and activated, we can assume that any observed effects were a direct result of the activity of this single enhancer and the interactions with the surrounding chromatin. One proposition is that the inserted element, when activated, acts as a switch to instruct the tissue-specific activation of downstream enhancer-like elements which themselves recruit cohesin. Given the lack of CTCF binding at these cryptic downstream sites within the newly formed domain, we speculate that the cohesin is most likely loading at these sites rather that stalling and accumulating as would be expected if there was a boundary element present on the chromatin.

The set of experimental results presented in this study bring us a step closer to understanding how 3D genome architecture can be directed via the recruitment of cohesin to active regulatory elements in a cell type specific manner. Once associated with the chromatin at these sites cohesin can spread by translocation, and via the loop extrusion mechanism, enable the formation of long-range chromatin contacts and domain formation. Peaks observed in ChIP-seq ultimately reflect the enrichment of cohesin at sites across the genome, therefore representing either loading or stalling of cohesin through the effect of boundary elements such as CTCF. Our set of experiments have not probed the dynamics of this mechanism as there is no way of pinpointing the exact site to which cohesin is initially recruited compared to where it translocates and stalls, an avenue that will be interesting to explore in future work.

## Materials and Methods

### CD34+ HSPC isolation and differentiation

CD34+ human stem and progenitor cells (HSPCs) were isolated from the three donors (001, 002 and 030), expanded and differentiated up until day 10 as previously descibed^53^.

### Mouse embryonic stem cell culture and genome engineering

E14TG2a.IV mESCs were maintained by standard published methods^54,55^. CRISPR/Cas9 mediated HDR strategies were used to produced genetically modified cell lines. mESCs were co-transfected with guide RNA and HDR vectors using Lipofectamine LTX reagent (ThermoFisher) according to manufacturer’s instructions. Sequences for guide RNA and HDR vectors are supplied in Table S2.

### Isolation of erythroid cells derived from adult mouse spleen

Mature primary Ter 119+ erythroblasts were obtained from spleens of female C57BL/6 mice treated with phenylhydrazine as previously described^56^.

### Embryoid body differentiation and erythroid population isolation

EB differentiation and CD71+ erythroid population isolation was performed following the previously published protocol^37^, aliquots were used fresh or frozen at -80°C depending on the downstream protocol.

### ATAC-seq

ATAC-seq was performed on 75,000 cells from target populations as previously described^34,57^. ATAC-seq libraries were sequenced with a high-Output v2 75 cycle kits on the Illumina NextSeq platform.

### ChIP-seq

ChIP-seq was performed on aliquots of 3-10×10^6^ CD71+ cells. For calibrated ChIP-seq, a 1% or 4% spike-in of WT HEK293 cells were added prior to sonication. Cells were double cross-linked using disuccinimidyl glutarate (DSG, Sigma) and 1% formaldehyde (Sigma) for a total fixation time of 1 hour. Fixed chromatin samples were fragmented using the Covaris sonicator (ME220) for 10 minutes (75 power, 1000 cycles per burst, 25% duty factor) at 4°C. 50µL per sample was removed as an input control. Sonicated samples were pre-cleared to remove background signal through incubation with a 1:1 mix of Protein A/G Dynabeads (InVitrogen). Antibody was added at a concentration of 1 µg/µL per sample and immunoprecipitated at 4°C overnight, a full list of antibodies used is supplied in Table S3. Immunoprecipitated samples were incubated with a 1:1 mix of Protein A/G Dynabeads for 5 hours at 4°C. Beads were then washed 4x in RIPA buffer on a magnetic stand, followed by 1x wash with TE (Sigma Aldridge) + 50mM NaCl (ThermoFisher). Chromatin was eluted from the beads using elution buffer and incubated for 30 minutes at 65°C with shaking. Input samples were diluted 1:1 with elution buffer. Samples and input controls were incubated at 65°C overnight for de-crosslinking, before RNase (Roche) and proteinase K (BioLabs) treatment. DNA fragments were purified using the Zymo ChIP DNA Clean & Concentrator kit (Zymo Research) and eluted in water. DNA concentration was quantified using the Qubit dsDNA HS assay (Invitrogen) as per manufacturers protocol. To assess sonication efficiency the D1000 Tapestation (Agilent) assay was performed on input sample. An approximately equal mass of input and IP DNA was used for indexing (0.5 ng – 1 µg). NEBNext Ultra II DNA Library Prep Kit (New England Biolabs) was used to prepare indexed sequencing libraries following the manufacturer supplied protocol. PCR amplification was performed for 7-11 cycles (depending on input DNA concentration) using NEBNext Mulitplex Oligos (New England Biolabs). Indexed sample concentration was quantified using the KAPA Library Quantification Complete Kit (Universal)(Roche). Samples were pooled as a 4 nM library and sequenced with a high-Output v2 75 cycle kits on the Illumina NextSeq platform.

### Next-generation Capture-C

Next-generation Capture-C was performed as previously described^56^ on 5×10^6^ mouse CD71+ cells derived from three separate differentiation experiments. Custom biotinylated DNA oligonucleotides used in Capture-C experiments are supplied in Table S1. Capture-C libraries were sequenced on the Illumina Nextseq platform using a 300-cycle paired-end kit (NextSeq 500/550 Mid Output Kit v2.5). To produce sufficient sequencing depth >1 x 10^6^ 150 bp paired-end reads were generated per viewpoint, per sample in multiplexed library.

### Tiled-C

Tiled-C experiments were performed on mESC and erythroid cells previously described^1,33^.

### RNA extraction

Total RNA was isolated from 1-5 × 10^6^ CD71+ from cultures of embryoid bodies. Cells were lysed in TRI reagent (Sigma-Aldrich) using a Direct-zol RNA MiniPrep kit (Zymo Research). DNase I treatment was performed on the column as recommended in the manufacturer’s instructions, with an increased incubation of 30 minutes at room temperature (rather than the recommended 15 minutes). RNA was eluted in DNase/RNase-free Water and degradation was assessed using RNA Screentape Analysis (Tapestation, Agilent Technologies) which determined the RNA integrity number (RIN) score of the sample. RNA was quantified either by Qubit RNA Broad-Range Assay (Invitrogen, ThermoFisher). RNA samples were stored at -80 °C prior to RNA-seq.

### Poly(A)+/- RNA-seq

1-2 µg of total RNA was depleted of rRNA and globin mRNA using the Globin-Zero Gold rRNA Removal Kit (Illumina) according to the manufacturer’s instructions. Samples were purified at -80°C overnight by ethanol precipitation, centrifuged for 1 hour at 12,000 g, washed twice with 70% ethanol, and resuspended in RNase-free water. RNA Screentape Analysis (Tapestation, Agilent Technologies) for quality control to ensure sufficient depletion and integrity of RNA samples. For mRNA enrichment: Poly(A)+ RNA was isolated, strand-specific cDNA synthesised, and the resulting libraries prepared for Illumina sequencing using the NEBNext Poly(A) mRNA Magnetic Isolation Module (New England Biolabs) and the NEBNext Ultra II Directional RNA Library Prep Kit for Illumina (New England Biolabs) following the manufacturer’s instructions. The poly(A)- RNA fraction was retained by storing the supernatant and all subsequent washes of the magnetic Oligo dT Beads bound with mRNA at -80 °C.

Poly(A)- samples were purified using Agencourt RNAClean XP beads (Beckman Coulter) and eluted in RNase free water. To remove any contaminating poly(A)+ RNA, an additional poly(A)+ selection was performed using the magnetic Oligo dT Beads from the NEBNext Poly(A) mRNA Magnetic Isolation Module following the manufacturer’s instructions. Poly(A)- RNA samples were purified using Agencourt RNAClean XP beads and eluted in the First Strand Synthesis Reaction Buffer and Random Primer Mix (2X) from the NEBNext Ultra II Directional RNA Library Prep Kit for Illumina. Fragmentation of RNA, strand-specific cDNA synthesis, and library preparation for Illumina sequencing were all performed using the NEBNext Ultra II Directional RNA Library Prep Kit for Illumina according to the manufacturer’s instructions.

For both Poly(A)+ and Poly(A)- samples: purification of the double-stranded cDNA, the adaptor ligation reaction, and the PCR reaction of adaptor ligated DNA were performed using Agencourt AMPure XP beads. Library profiles were assessed by visualisation on a D1000 tape on a Tapestation (Agilent Technologies) and quantified using a universal library quantification kit (KAPA Biosystems, Roche: 07960140001). Poly(A)+ and poly(A)- RNA-seq libraries were sequenced on the Illumina Nextseq platform using a 75-cycle paired-end kit (NextSeq 500/550 High Output Kit v2.5: 20024906).

### Data Analysis

#### ATAC-seq and ChIP-seq

For publicly available data, raw data was downloaded in SRA format from the relevant GEO repositories. Using the SRA Toolkit^58^ ‘fastq-dump’ function, fastq files were extracted. ATAC-seq and ChIP-seq data in fastq format were analysed using the CATCH-UP pipeline^59^. In brief, single- or paired-end reads were aligned to the target genome using Bowtie2^60^. Samtools^61^ was used to remove duplicates and index bamfiles. A coverage track of the aligned reads was generated using deepTools^62^ ‘bamCoverage’, where coverage was calculated for a single base pair window and replicates were normalised to Reads Per Kilobase per Million mapped reads (RPKM) using flags ‘-bs 1--normalizeUsing RPKM’. The output coverage files in bigWig format were visualised using the UCSC Genome Browser^63^.

#### Calibrated ChIP-seq

The pipeline for the analysis of calibrated ChIP-seq data was adapted from the scripts used in Fursova et al.^64^, and is available in the UpStreamPipeline GitHub repository. In brief: (1) whole genome fasta files for the target and spike-in species were concatenated and used to build Bowtie2 index files; (2) paired-end reads were then aligned to the concatenated genome using Bowtie2^60^; (3) reads were separated and counted based on alignment to target or spike-in genome and a down-sampling factor was calculated based on the total number of spike-in reads per sample as follows:

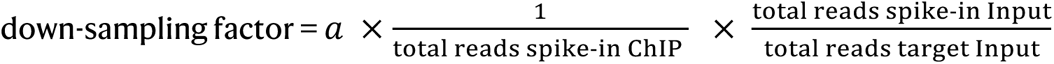

where *a* = coefficient for bulk normalisation of samples within a single experiment to enable the value of the largest down-sampling factor to equal 1^64^. (4) using the calculated down-sampling factor, reads for each ChIP sample were randomly subsampled using Sambamba^65^. (5) Input samples were used to correct the ratio of spike-in to target read counts across the biological replicates to account for minor variations in the mixing of spike-in cells, (6) sub-sampled bamfiles were then processed according to the standard ChIP-seq analysis method described above.

#### Next-generation Capture-C

Capture-C data were analysed as previously described^56^ aligning to the mouse mm9 reference genome. CaptureCompare was used to generate comparisons of the Capture-C interaction profiles (https://github.com/djdownes/CaptureCompare). Briefly, unique interactions were normalised to cis interactions, averaged (n = 3), and the difference between the means calculated. For visualisation, means and standard deviations were binned into 150 bp bins and smoothed with a sliding window of 3 kb, and plotted using ggPlot in R.

#### Tiled-C

Tiled-C data was analysed using the tCaptureC workflow available in the UpStreamPipeline GitHub repository. This uses Hi-C Pro^66^, with DpnII digestion of the genome using the built-in ‘digest-genome’ utility. Capture target was specified using an input BED file containing region coordinates, min/max fragment sizes were specified in the config file as 20 and 100,000 respectively, and ICE normalisation implemented. All other parameters applied were set at default. HiCPlotter^67^ was used to visualise the interaction matrices.

#### Custom Genome Builder

To correctly align reads to the genetically engineered R2-insertion cell lines a custom genome was built. In brief, this pipeline takes a fasta file containing the insert sequence and a BED file of the coordinates of the edited region, and bioinformatically cuts-and-pastes together a custom genome. Samtools^61^ was used to index the custom genome fasta file, and Bowtie2^60^ used to create new indices.

#### Poly(A)+/- RNA-seq

RNA-seq reads were aligned to the genome (mm39) using STAR^68^ with --outFilterMultimapNmax value set to 2. Alignment reads were filtered for proper pairs, sorted, and PCR duplicates removed using Samtools^61^. Biological replicates were normalised to RPKM in a strand-specific manner using deepTools^62^ bamCoverage and the flag--filterRNAstrand. Biological replicates were merged and converted into bigWig file format for visualisation in the UCSC genome browser.

#### Allelic skew and meta analysis

Phased genomes were variant called on 10x sequencing data using longranger (v2.2.2). ATAC-seq read counts for skew analysis were obtained using pysam (v0.20.0) and matched by haplotype. Variants were identified when found on all haplotypes with a positive, or on all haplotypes with a negative skew. Only variants within the peak called regions were analysed.

51 regions across the genome were selected based on skew analysis in which the three donors were a mix of homozygous-ref, heterozygous and homozygous-alt where the alternative haplotype was shown to exhibit differential signal (or skew) in ATAC-seq, with stringent p-value threshold of p < 0.0001. To calculate the RAD21 coverage across the 51 regions, the BedTools^69^ ‘multicov’ function was used, with a bedfile of the 51 regions and bamfiles of RPKM normalised RAD21 ChIP-seq data from the three donors. Data was visualised using the Python (3.11.3) package Seaborn (0.12.2), with p-values calculated using the Scipy (1.10.1) paired T-test.

#### RAD21 coverage analysis for enhancer KO models

Two regions: (1) chr11: 32188452 – 32249902 and (2) chr11: 32323049 – 32384895 were provided in bed format along with bamfiles of RAD21 ChIP-seq for WT, R2-only and SE KO models, and to calculate coverage scores for each replicate using the BedTools^69^ ‘multicov’ function. Data was visualised using the Python (v3.11.3) package Seaborn (v0.12.2).

#### CTCF motif analysis

CTCF ChIP-seq data was peak called using Lanceotron^70^ with default settings and fasta sequence for each peak extracted using the Bedtools ‘getfasta’ utility^69^. The MEME frequency matrix for CTCF (CTCF_MA0139.1.meme) was downloaded from JASPAR^71,72^. Using the FIMO (Find Individual Motif Occurrences)^73^ tool from The MEME suite^71^, occurrences of MA0139.1 were determined within the CTCF peak calls using default parameters. Peaks with identified CTCF motifs were annotated with the orientation.

### Data availability

ATAC-seq, ChIP-seq, RNA-seq, Capture-C and Tiled-C data generated for this study have been deposited in the Gene Expression Omnibus (GEO) under accession code GSE244929. Previously published ChIP-seq and ATAC-seq data that were reanalysed here are available under the following accession codes: GSE57092, GSE30203, GSE30203, GSE36028 and GSE97871. Previously published Tiled-C data that were reanalysed here are available under the accession code GSE137477. Previously published data from the R2 only and DSE models that were reanalysed here are available under the accession code GSE220463.

All other data supporting the findings of this study are available from the corresponding author on reasonable request.

### Code availability

Pipelines for ATAC-seq, ChIP-seq, Calibrated ChIP-seq, RNA-seq, Tiled-C and building the custom genomes are all available in the following GitHub repository: https://github.com/Genome-Function-Initiative-Oxford/UpStreamPipeline. Version numbers for each tool used within the pipelines are provided in the upstream.yml file.

## Acknowledgements

The authors would like to express their gratitude to all colleagues who contributed to this work, in particular: of the Weatherall Institute of Molecular Medicine genome engineering facility, and the Higgs lab for providing the R2-only and DSE cell lines. This work was supported by a Wellcome Strategic Award (106130/Z/14/Z to J.R.H.), and the Medical Research Council (MRC) (MC_UU_00016/14 to J.R.H., 4050189188 to D.R.H.). E.G., C.L.H, H.F., and J.B. were supported by Wellcome Doctoral Programmes (108861/Z/15/Z, 109110/Z/15/Z, 109097/Z/15/Z, and 219979/Z/19/Z). T.A.M. was supported by Medical Research Council (MRC, UK) Molecular Haematology Unit grant MC_UU_00016/6 and MC_UU_00029/6.

## Author contributions

The authors contributed as follows: Conceptualisation of project: E.G., C.H., M.K., J.H., D.H., cell line generation: C.H., N.R., H.S.F., M.K, NG Capture-C experiments: C.H., Tiled-C experiments: A.M.O, J.B., ChIP-seq experiments: E.G., C.H., bioinformatic analysis: E.G., S.G.R., E.S., C.H., project supervision: J.H., M.K., T.A.M., D.R.H., writing of manuscript: E.G., review and editing of manuscript: D.H., J.H.

