## Supplementary Material for "Active regulatory elements recruit cohesin to establish cell-specific chromatin domains"

**Fig. S1: Natural variation example at chr6:79,507,436-79,722,535.**

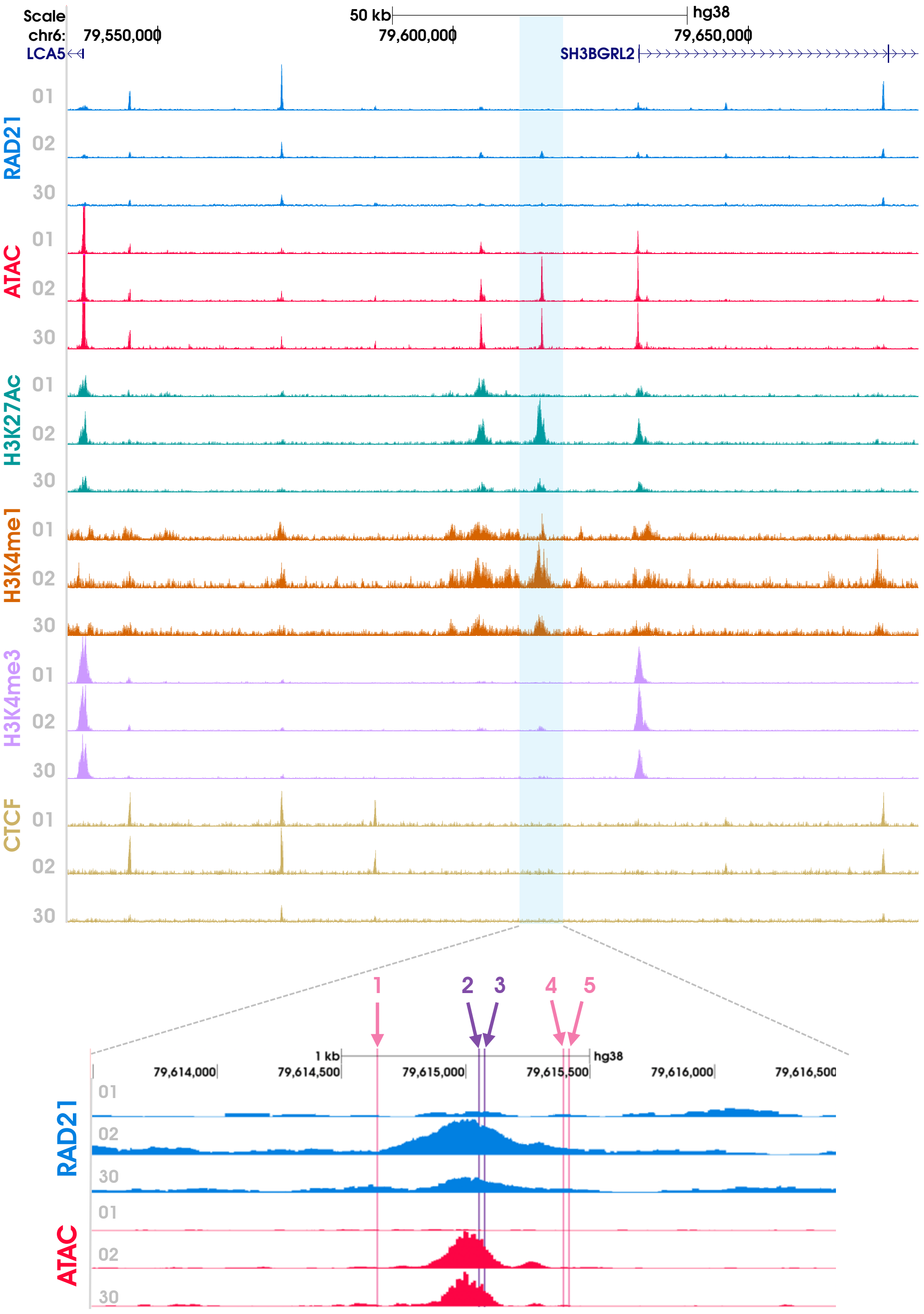


| | Position | Don001 | Don002 | Don030 |
|---|---|---|---|---|
| 1 | chr6:79,614,643 | G | G > A | G > A |
| 2 | chr6: 79,615,053 | A > AC | A | A |
| 3 | chr6: 79,615,075 | A > G | A | A |
| 4 | chr6: 79,615,393 | A | A > C | A > C |
| 5 | chr6: 79,615,415 | C | C > T | C > T |

reference = peak

reference = no peak

Region shown is chr6:79,507,436-79,722,535 (hg38). Numbers 01, 02, 30 indicate the anonymized donor identifiers. ChIP-seq for RAD21, CTCF and histone marks (H3K4me1, H3K4me3, H3K27ac) are shown for each donor in the top panel along with the open chromatin signal (ATAC-seq). The region highlighted in blue is shown in detail below. Here the difference in signal across the donors can be clearly seen: donor 1 is homozygous and displays a positive ATAC-seq signal and the greatest peak of RAD21, donor 2 is heterozygous and displays a positive ATAC-seq signal and a moderate peak of RAD21, donor 30 is homozygous with no ATAC-seq or RAD21 signal. The pink line indicates the SNP we have identified as potential causal for this difference across the donors.

### Fig. S2: Natural variation example at chr1:154,364,808-154,566,107.

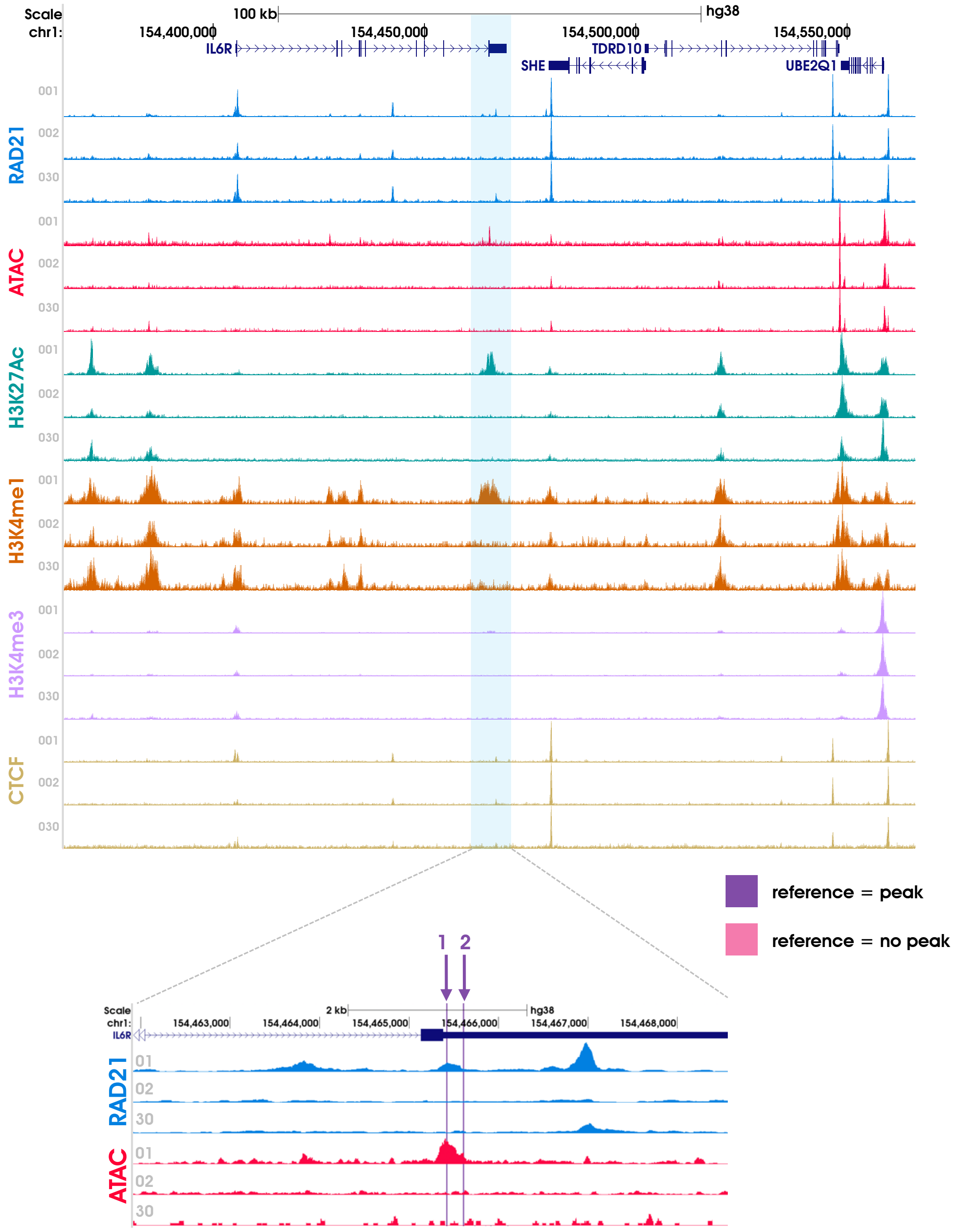


| | Position | Don001 | Don002 | Don030 |
|---|---|---|---|---|
| 1 | chr1:154,465,420 | ref. T | alt. T>C | alt. T>C |
| 2 | chr1:154,465,608 | ref. C | alt. C>T | alt. C>T |

Region shown is chr1:154,364,808-154,566,107 (hg38). Numbers 01, 02, 30 indicate the anonymized donor identifiers. ChIP-seq for RAD21, CTCF and histone marks (H3K4me1, H3K4me3, H3K27ac) are shown for each donor in the top panel along with the open chromatin signal (ATAC-seq). The region highlighted in blue is shown in detail below. Here the difference in signal across the donors can be clearly seen: donor 1 is homozygous and displays a positive ATAC-seq signal and the greatest peak of RAD21, donor 2 is heterozygous and displays a positive ATAC-seq signal and a moderate peak of RAD21, donor 30 is homozygous with no ATAC-seq or RAD21 signal. The pink line indicates the SNP we have identified as potential causal for this difference across the donors.

**Fig. S3: Comparison of the ATAC-seq reads originating from the R2 enhancer in WT versus the R2-insertion erythroid cells.**

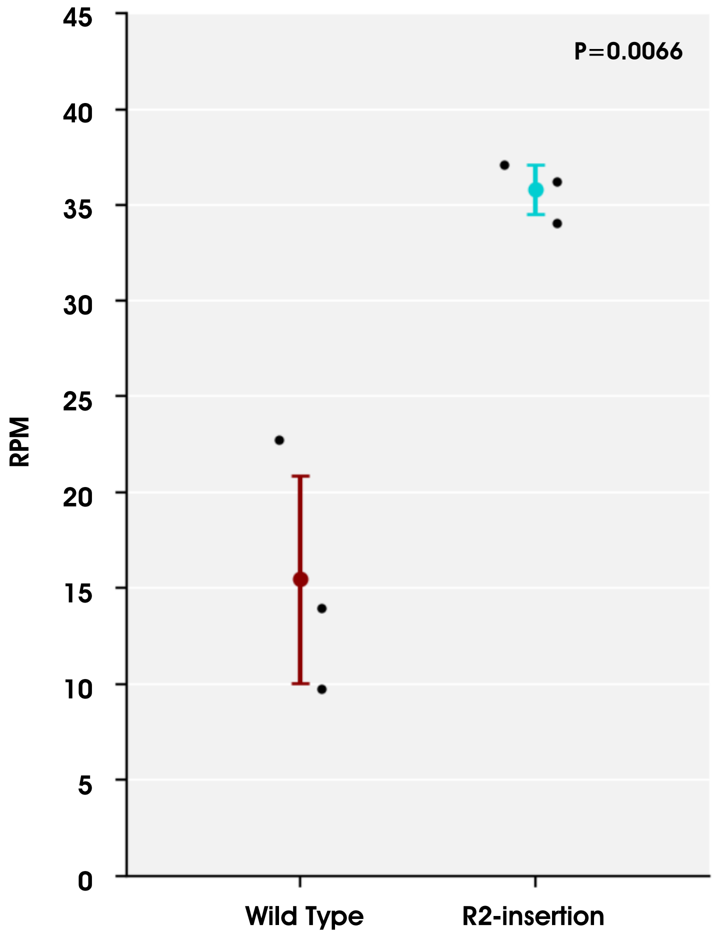


Reads that overlapped with the 325 bp R2 sequence normalised to reads per million (RPM) for ATAC-seq performed on WT and R2-insertion erythroid cells. Mean and standard deviations for each model are shown, each point represents a biological replicate/independently targeted clone. The mean of R2 reads from ATAC-seq in the R2-inseryion erythroid cells significantly differ from that of WT, p-value (p=0.0066) shown was calculated using a two-tailed student's T-test.

**Fig. S4: Identification of an activatable region devoid of active histone marks, open chromatin and CTCF binding.**

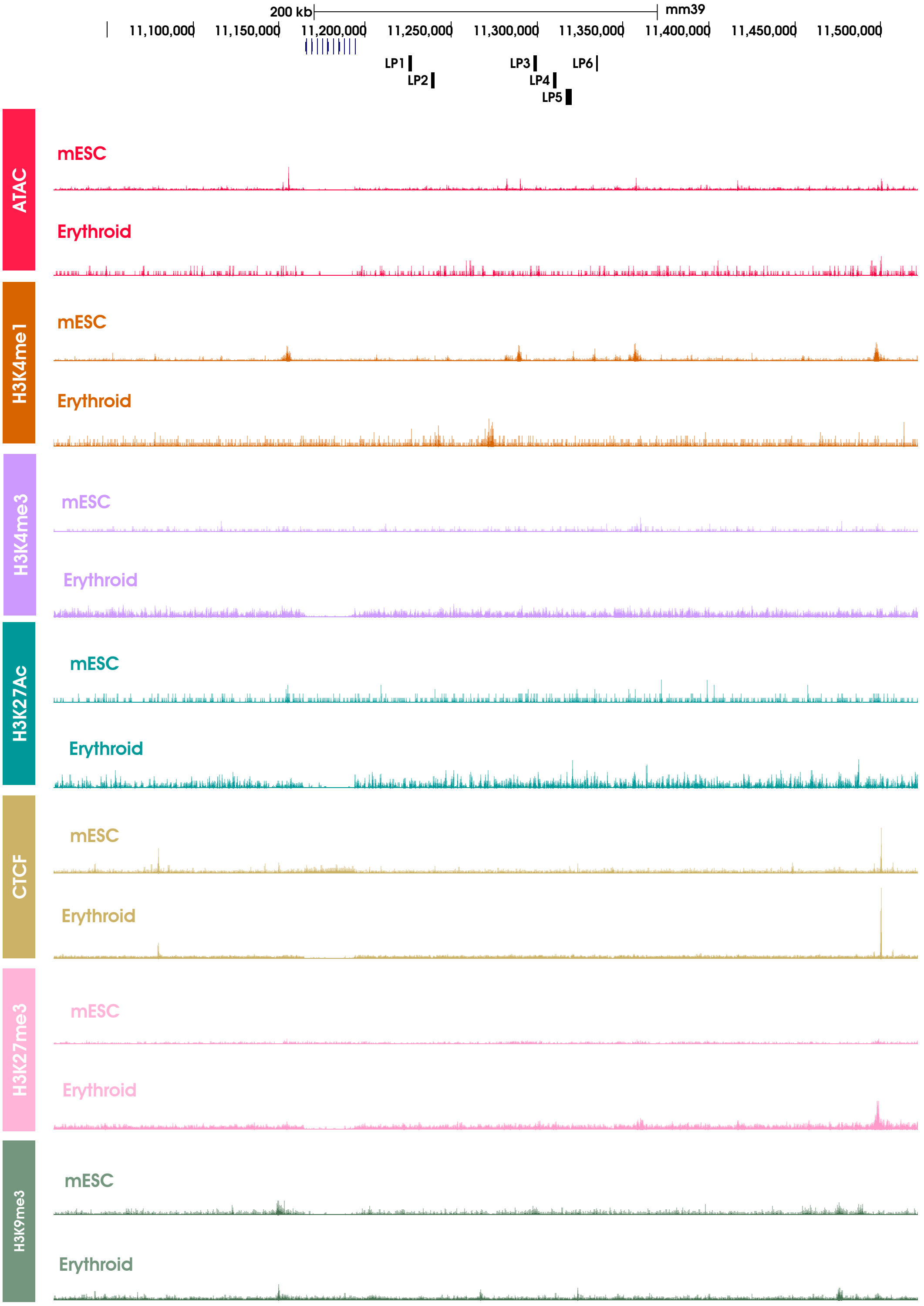


Characterisation of the chromatin environment across chromosome X was performed and the region chrX:11,018,725-11,521,742 (mm39) was identified to fit all the criteria to enable targeting of R2 to this site.   All data is RPKM normalised.  Publicly available data sources are listed in Supplementary Table 4, all other data is available to download from GSE244929.

**Fig. S5: There is no underlying 3D structure within the selected chromosome X region.**

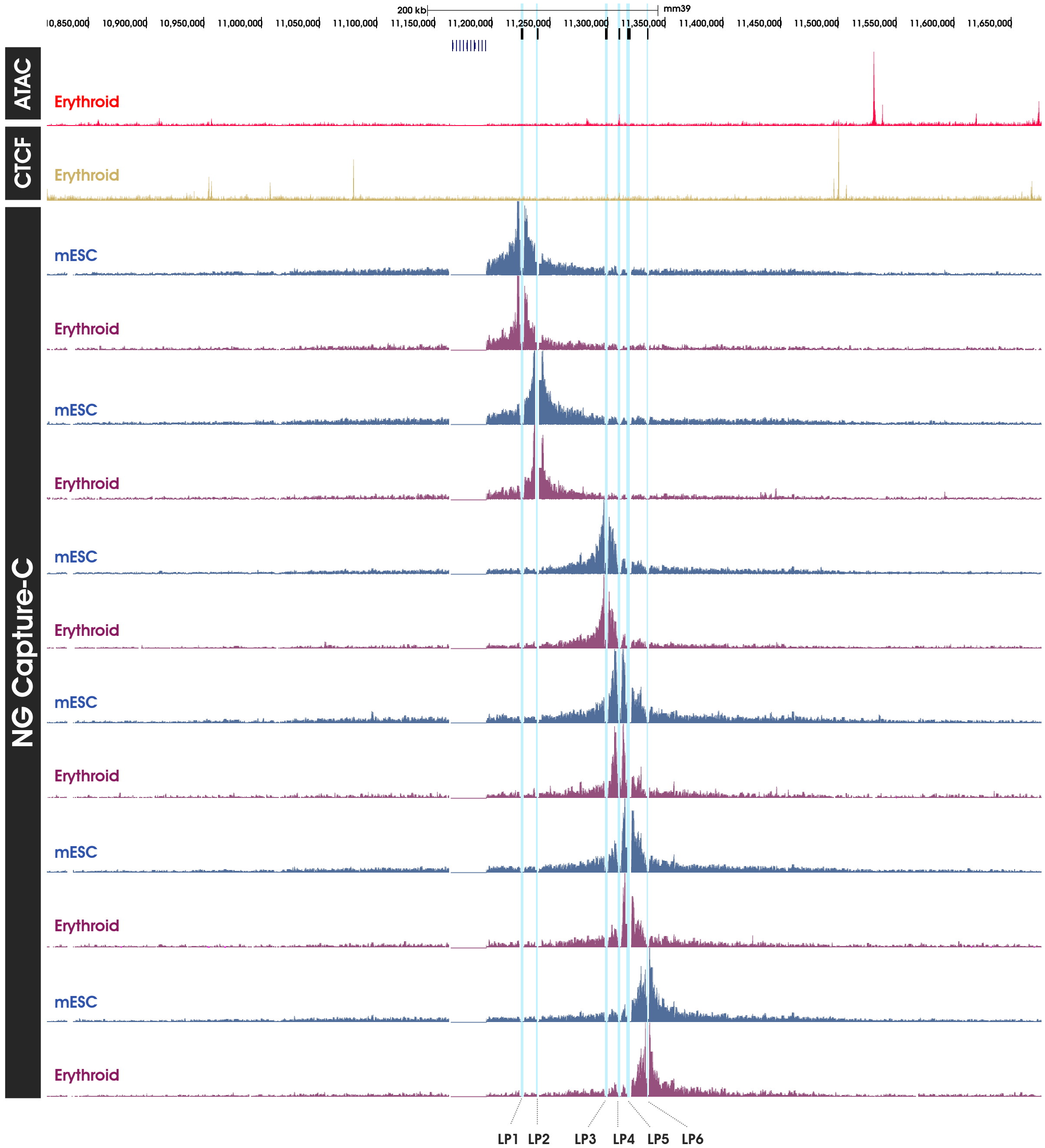


Next-generation (NG) Capture-C was performed from the six viewpoints shown (LP1-LP6) and highlighted in blue. Data from WT mESC is shown in blue and WT erythroid cells in purple. RPKM normalised ATAC-seq (dark pink) and CTCF (gold) data in WT erythroid cells are shown for reference.

**Fig. S6: Characterisation of the histone landscape in WT cells.**

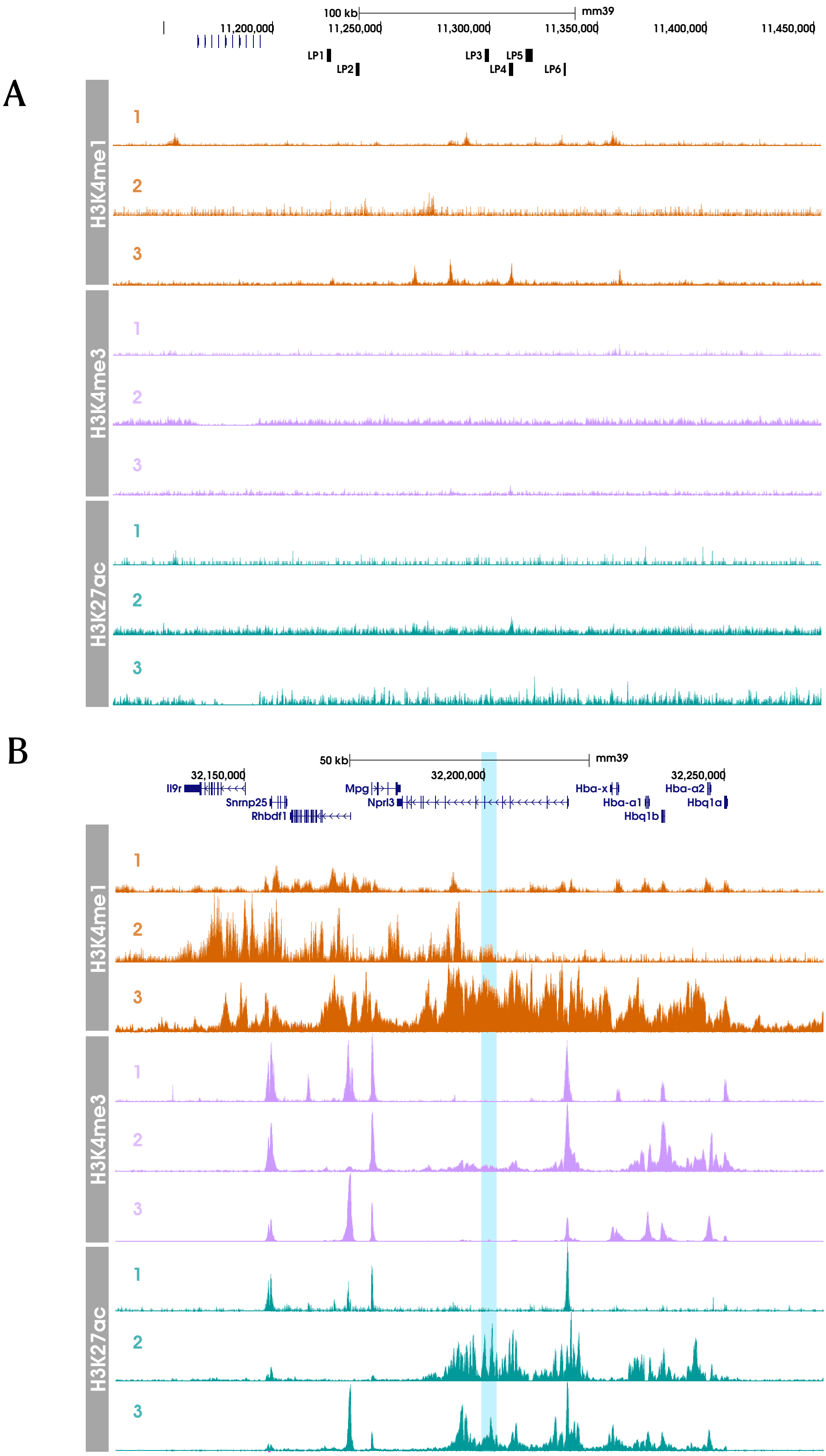

The histone modifications H3K4me1 (orange), H3K4me3 (lilac), H3K27ac (turquoise) are compared across three unedited WT mouse cell types: ESCs (no. 1), primary erythroid cells derived from spleen (no. 2) and erythroid cells derived from embryoid body cultures (no. 3), across two loci: (A) the chromosome X region (chrX:11,126,444-11,453,282) that was selected for targeting, and (B) the alpha globin locus (chr11:32,123,214-32,270,752). The location of the endogenous R2 enhancer in the alpha globin locus is highlighted in blue for reference. Data sources for publicly available data sets are detailed in Supplementary Table 4, all other data was generated by the authors and is available in GSE244929.

**Fig. S7: Characterisation of the histone landscape in WT and R2-insertion embryoid body derived erythroid cells.**

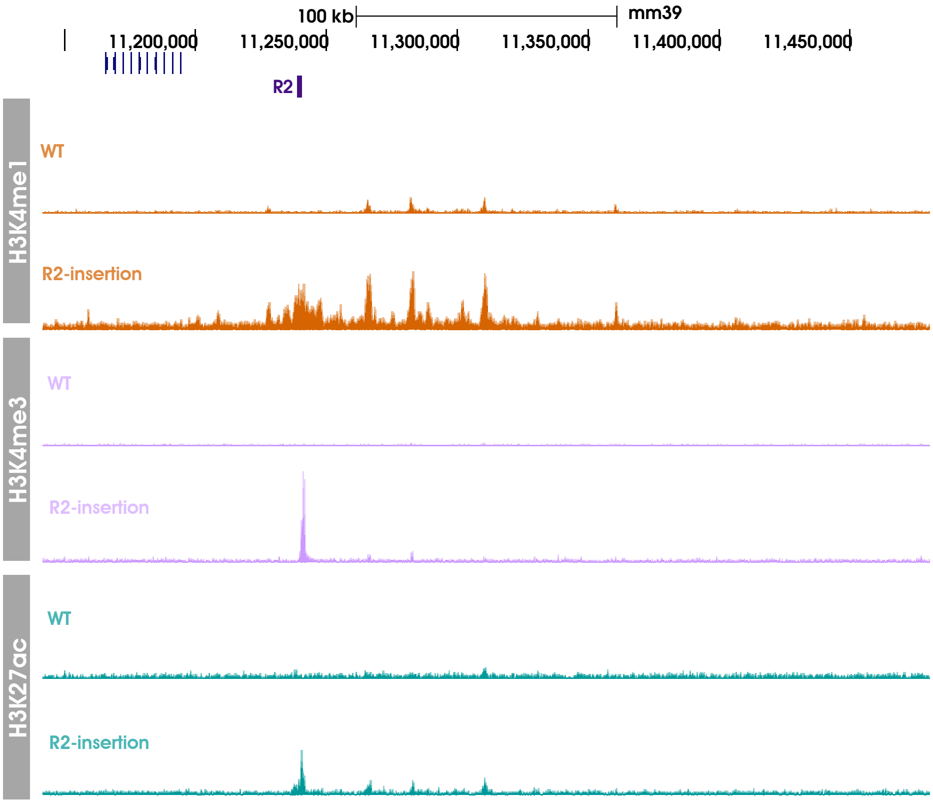


The histone modifications are shown as follows: H3K4me1 (orange), H3K4me3 (lilac) and H3K27ac (turquoise), across the region chrX:11,126,444-11,453,282. The R2-insertion site is indicated above the tracks by a purple rectangle. All tracks are RPKM normalised. Raw and processed data are available to download from GSE244929.

**Table S1: Capture oligonucleotides were designed to six viewpoints (LP1-LP6) across the chromosome X locus.**

| Viewpoint | Capture DNA oligo | Sequence (5′-3′) |
|---|---|---|
| LP1 | 5′ | [Btn]GATCTATATGGGATGAGATATTAGTTGGGGTTCTCCAGAGACAGGGAGC<br>CAATAAGATGTATGTTGTTGG |
| | 3′ | [Btn]CCCAGCCAAGAGAATGGTGTCATTCACAACTGGCTGGGTTTTCTTGTAT<br>CAATTAAGAGTCAAGATGATC |
| LP2 | 5′ | [Btn]GATCATGGAATATTTTTTCTTTTGGACTTGCTTCACTCAAATAAAAAAAC<br>TGTTCACCTCATGTTTTCCA |
| | 3′ | [Btn]TGAATTCCCTGAAGGTAAGAACAGGAATTATCTTATCTCTAAACCAGGT<br>ATTGGGCAATTCAGGTGGATC |
| LP3 | 5′ | [Btn]GATCATTAAAGGAGACAGGAGGAAAGGATACCATCCCCGCAGCAGGGA<br>AATGGACTGCAACAGAGCATCC |
| | 3′ | [Btn]ATGAGGCTTTGTCCTAAGCGGCTAAGTGTCCCCGGTGTTCCTAGCAGGA<br>ATACAGGCTGGCTCATGGATC |
| LP4 | 5′ | [Btn]GATCCAAAATGCTCTGCTGAACTCAAGATAGTTCATTTCACTAAAGGCC<br>GTGCTGAGGAGCAGGGGCAAA |
| | 3′ | [Btn]GCTACCATGTAAGCCCCAGGACCAGCTATGTCTTGCTCCTGGAGAAGGC<br>GGTCATCATGGCTACCTGATC |
| LP5 | 5′ | [Btn]GATCTAGTCCTCACACATTTATGAGATGGTGTCACCAAGAGCACCTTCA<br>CAAGAAGACCAATGCGATACC |
| | 3′ | [Btn]TACTTTTCAAAGAAACTAAAAATTCTCACTACCTGCATTCATCATCCAAA<br>ATTCTCTAGAGAAACAGATC |
| LP6 | 5′ | [Btn]GATCCAAATGTTCTTAAGGTCCTGAAGTAGTTTTTAAAAATAGACTTCCG<br>TGAAGAAAGAACTCTGGTTG |
| | 3′ | [Btn]TATTTTAAAAGAAGAAAAGGAATAGACATTTGTGTAGGTCATGTTGCTG<br>GAGTCTAGAAAGCCTAAGATC |

**Table S2: Sequences for guide RNA and HDR vectors.**

| Target | Sequence 5′-3′ |
|---|---|
| LP2 guide | GGAGAGTAGTGGCCCAACTCT |
| LP2 R2-insertion donor | CCCACAATGGCATGGCATTGCTGAATGTATCCCAGTTTTTTGGCTTTTGATGCCTATATTGTCAGGG<br>TCATATATCCCCAACCACTACTGAGAAAAACTTCCTTGGGTCAGTGTTGGATAGCAGCAGCTCCCTT<br>ATGCTTACTTGGAGTGGTTCTAAGATTTTGTGTTTTACATTAGGTATTTTATCTATTTGCAGTTGGTT<br>TGGGATGTTGTGTGAGACAAGGGTCCAGTTTCTTTCTTCTGCATGTACACATCTAGTCTTCAAACAG<br>AGTTCAATGAAGAAATTGTCAATTTTCTATTGTATATACTTTTGCCATATTCATGCTGGAAATGGCT<br>TTCTTTGCTTTTCCCAAGGTAATCTGCATTTAATGTGGATGAGAGGATGCTGGCCATAACCCCCCCC<br>CCCTTCCATCCCTGCACTGGAAAATATTCTGGGCATCTGACAGTCTATAAGACCATGCTCTTCCCAG<br>GTAGTTATCTTCTTATGACTGAGCACAGTTGGTTGATCTACTGCATACCTACACTAACTAGGTCAAA<br>GTAGCATACACCCATCTGGAACCTATCAGTGACCATAGTCAACAGCAGGTGTACACACCCAGGCCA<br>AGGGTGGAGCAGACCACTGTGGGATCTATGGAGATGCTTGAACGAGCAGATAACTAAGCCAAGCAT<br>GACTCAGAGTTTCTAGAGGCCACTAGGACTGCTGAGTAATACTTGGGGGTACAGAGTCAGAAAGGA<br>AAGGACAAATGGTACCACTGATTAGGACCTCTGACGCTGTTTTCCCATCCTGTTTATTTGCCAAGTG<br>ACCCTGTGCCTGTTACCTTAAGTCAAAAACTTATGACTTTTCTGATGGGCAGTTTTTCCCCTGGTCT<br>CCACTGGCCTGAAGAGACGTTTTCAAAGTGATGCTTTTCAGCTGAGGTGT<br>ATTCCATCCTTCCCTCTGCTTCCACAGGTCTTGGATGGACAACCTCTGTTTGTTCCATTTATCTTTT<br>TGCAGGTTTCTAGCCCCATCTAGGTGTCTGCTTCCAGGAGGACAAGAACTGGCATACATAACCATA<br>AAGTCTCAACAACGCAGTAGGTGTTTTCCAGATATCCTCCCCTGAGTTCTTTCCAGCTGAATGAGG<br>GAGCCACTGAGGTCTGTGCCTTTGCTTACCTCCCCAACTACCATGCAGCAGCCTTGACAACAGCAC<br>TTCTGTTCTGCTATGCAAATTTGTTTTCTCTCTCTACTGAATTCCCTGAAGGTAAGAACAGGAATTA<br>TCTTATCTCTAAACCAGGTATTGGGCAATTCAGGTGGATCATGCATGATACTCTGGGTCACACATCC<br>CCC |
| RAD21 guide | CCTCAGATAATATG GAACCGTGG |
| RAD21$^{TST}$ donor | CCCGGGTTGGAAGGTTATCAGGGGCCGGTTTGATTTTGGTTTTGTTTTCACTTTAAAATCTGCTGAG<br>TGTTTGTTTTTGCTAACTCACATCTCTGTTGTGGCGCCTTCACTTACTTCTTGAACTCTGTATGCCT<br>TAAGATAATATGCTTTAGATAATAAGGCCTTTACCTTAGTGACTAGCATGGGAACCACTTGCGTAAC<br>TACACGAATGTCCAGTCCCAGCTCCAGCCAAACGGAGCTCACTTTGACCAGTGCCAAAATTGCATC<br>TTCTGGTTACTACTTTGTGCGTGAGTTACTTGAAATCATCTGCTTTGTTTTGTTCTATTTCAGCGAG<br>CTCTTGCTAAAACTGGAGCAGAGTCTATCAGTTTGCTTGAGCTGTGTCGAAACACAAACCGAAAGC<br>AGGCAGCAGCAAAGTTCTACAGCTTTTTGGTTCTTAAGAAGCAGCAAGCCATCGAGCTCACACAGG<br>AAGAGCCGTACAGTGACATCATTGCAACGCCCGGCCCACGCTTTCACATCATCAGCGCCTGGAGCC<br>ACCCCCAGTTCGAGAAGGGCGGCGGCAGCGGCGGCGGCAGCGGCGGCAGCGCCTGGAGCCACCCC<br>CAGTTCGAGAAGTGAGATATCGGAGCTAGATGTGTTCGAGCTAGTGATAACTCACTAGTACATACA<br>AATTGCCCCCGTGTGCAGGGCACCAAAACCCTTTAAGAAAGTTTTTAGATTTCTGTTTGTACAAAAA<br>TCTTTGCCTTTTCTTTCTTCTTTTTCCCCCCAGTGTTTCTAATTTTGTCAACCATATTTTTAAGGGAA<br>ACTGCTTATTTGGGTTGGGTTTGTATTCCTGGAGAAAACAGTAGCCCAAGAACCCAGAAGACTTTTA<br>ACAGTTCAGAACAGATGTGTGCAATATTGGTGCATGTAAGAATATGGAGTAACAGTCAAAAGGCAC<br>CATTTTTAATGTTAGTTTTCCATTACTATGTTGAAAGGAAAACCTGCCTAGGAAAATGCCTGACACT<br>TTAAGAACTGTGGTTTGAGTCCCTTGACAGGAAGAGAAAAATGTCTTCCCATCAGTGAAACCAACG<br>GTCTGGTTAACCACTGTAGTAGGGATAGTGTGTGAAGCATCCCGGG |

**Table S3: Antibodies used for ChIP-seq experiments.**

| Target | Supplier | Catalogue # | Lot # |
|---|---|---|---|
| StrepTag | Qiagen | 34850 | - |
| CTCF | Merck Millipore | 07-729 | 2836926 |
| H3K4me1 | abcam | ab195391 | |
| H3K4me1 | abcam | ab8895 | GR3206285-1 |
| H3K4me3 | Merck Millipore | 07-473 | |
| H3K4me3 | abcam | ab8580 | GR3190162-1 |
| H3K27Ac | abcam | ab4729 | GB3205523-1 |
| RAD21 | abcam | ab1546769 | 1035529-6, 1035529-4 |
| RAD21 | abcam | ab992 | GR3310168-13 |

Table lists the antibodies used in the ChIP-seq experiments in this study, and lot numbers are provided if known. Specific information for each experiment is provided in the GEO submission (GSE244929).
